# Medium-chain acyl-CoA dehydrogenase, a gatekeeper of mitochondrial function in glioblastoma multiforme

**DOI:** 10.1101/2020.09.28.316554

**Authors:** Francesca Puca, Fei Yu, Caterina Bertolacci, Piergiorgio Pettazzoni, Alessandro Carugo, Emmet Huang-Hobbs, Jintan Liu, Ciro Zanca, Federica Carbone, Edoardo Del Poggetto, Joy Gumin, Pushan Dasgupta, Sahil Seth, Frederick F. Lang, Erik Sulman, Philip L. Lorenzi, Lin Tan, Mengrou Shan, Zachary P. Tolstyka, Maureen Kachman, Li Zhang, Angela K. Deem, Giannicola Genovese, Pier Paolo Scaglioni, Costas A. Lyssiotis, Andrea Viale, Giulio F. Draetta

## Abstract

Glioblastoma (GBM) is among the deadliest of human cancers. Despite extensive efforts, it has proven to be highly resistant to chemo- and immune-based therapeutic strategies, and little headway has been made with targeted inhibitors. Like many cancers, metabolism is dysregulated in GBM. Thus, to identify new vulnerabilities and drug targets in GBM, we conducted genetic screens using pooled RNAi libraries targeting metabolic enzymes. We screened multiple glioma stem cell-derived (GSC) xenograft models, which revealed that several enzymes involved in the mitochondrial metabolism of fatty acids were required for tumor cell proliferation. From among these, we focused on medium-chain acyl-CoA dehydrogenase (MCAD), which oxidizes medium-chain fatty acids, due to its consistently high score across all of our screens, as well as its high expression level in multiple GSC models and its upregulation in GBM compared to normal brain.

In this manuscript, we describe the dependence of GBM on sustained fatty acid metabolism to actively catabolize lipid species that would otherwise damage the mitochondrial structure. The uptake of medium-chain fatty acids lacks negative feedback regulation; therefore, in the absence of MCAD, medium-chain fatty acids accumulate to toxic levels, inducing reactive oxygen species (ROS), mitochondrial damage and failure, and apoptosis. Taken together, our findings uncover a previously unappreciated protective role exerted by MCAD in GBM cells, making it a unique and therapeutically exploitable vulnerability.

## INTRODUCTION

Glioblastoma multiforme (GBM) has a median survival of 15 to 19 months (Liebelt et al., 2016; Ru et al., 2013; Stupp et al., 2005). Despite extensive efforts, studies of both signal transduction inhibitors and immune-targeted agents have failed to improve the prognosis for patients with these tumors (Gan et al., 2017; Hou et al., 2006; Reifenberger et al., 2017; Weller et al., 2015; Yang et al., 2017). As with most cancers, altered metabolism is a defining characteristic of glioblastoma. Dependency on aerobic glycolysis (Warburg effect), as well as on glutaminolysis, are well documented (DeBerardinis et al., 2007; Portais et al., 1996; Warburg, 1956), and recent landmark studies have identified mutations in GBM that directly impact mitochondrial metabolism (2008; Brennan et al., 2013). However, the molecular drivers underlying this metabolic reprogramming are only partly understood, and therapeutically actionable metabolic dependencies have not been fully determined. In the present study, we applied *an-in vivo* genetic screen platform (Carugo et al., 2016) to orthotopically implanted, patient-derived 3D glioma tumor spheres and identified medium-chain acyl CoA dehydrogenase (MCAD), a mitochondrial enzyme that catalyzes the first step of medium-chain fatty acid (MCFA) β-oxidation (FAO), as a node of metabolic vulnerability for GBM tumor growth and maintenance. Targeting MCAD in primary GBM models yielded catastrophic levels of cell death triggered by massive accumulation of unmetabolized MCFAs, which induced irreversible mitochondrial alterations, oxidative stress and lipid peroxidation in tumor cells. Thus, in GBM, the activity of MCAD does not solely support cellular energetics as part of the β-oxidation cycle, but our work demonstrates a previously unappreciated protective role for MCAD fatty acid catabolism to clear toxic accumulations of lipid molecules that may otherwise cause lethal damage to the cell. Given that inherited MCAD deficiency is a condition compatible with normal quality of life, with individuals carrying genetic bi-allelic inactivation of its encoding gene (*ACADM*) requiring only dietary adjustment to thrive (Schatz and Ensenauer, 2010; Vishwanath, 2016), an actionable therapeutic opportunity exists to target MCAD in GBM patients.

## RESULTS AND DISCUSSION

To achieve a functional output of metabolic gene functions that might be essential for GBM cell survival, we generated a prioritized list of 330 metabolism genes and created a barcoded, deep-coverage (10 shRNAs/gene) shRNA library that encompasses a wide range of metabolic pathways and activities (Figure S1A). As a disease model, we selected low-passage, 3D-cultured, patient-derived glioma tumor-initiating cells (GSCs) established at the MD Anderson Brain Tumor Center (Hossain et al., 2015; Lang et al., 2018; Thomas et al., 2018). Among over 70 genomically characterized GSCs available to us, we selected two carrying distinct gene expression signatures classified as belonging to the proneural (GSC 8.11) or classical (GSC 6.27) (Figure S1B,C) subtypes. After shRNA library transduction, cells were implanted intracranially, and tumors were grown until mice showed tumor-related neurological symptoms. At endpoint, tumors were excised and processed as previously described (Carugo et al., 2016) (Figure 1A). Analysis of the most significantly depleted shRNAs in both GBM models uncovered the potential role of several mitochondrial enzymes involved in fatty acid metabolism in GBM growth (Figure 1B; Figure S1D-F). Given compelling evidence of its role in fatty acid metabolism in brain tissues, we focused our attention on MCAD, which oxidizes medium-chain (4- to 12-carbon) fatty acids (MCFAs). MCFAs such as octanoate are normally present in plasma at significant levels (Mamunes et al., 1974) and readily cross the blood-brain barrier (Spector, 1988). It has been shown that up to 20% of brain oxidative metabolism can be attributed to octanoate use in an intact rodent physiological system (Ebert et al., 2003).

**Figure 1.**
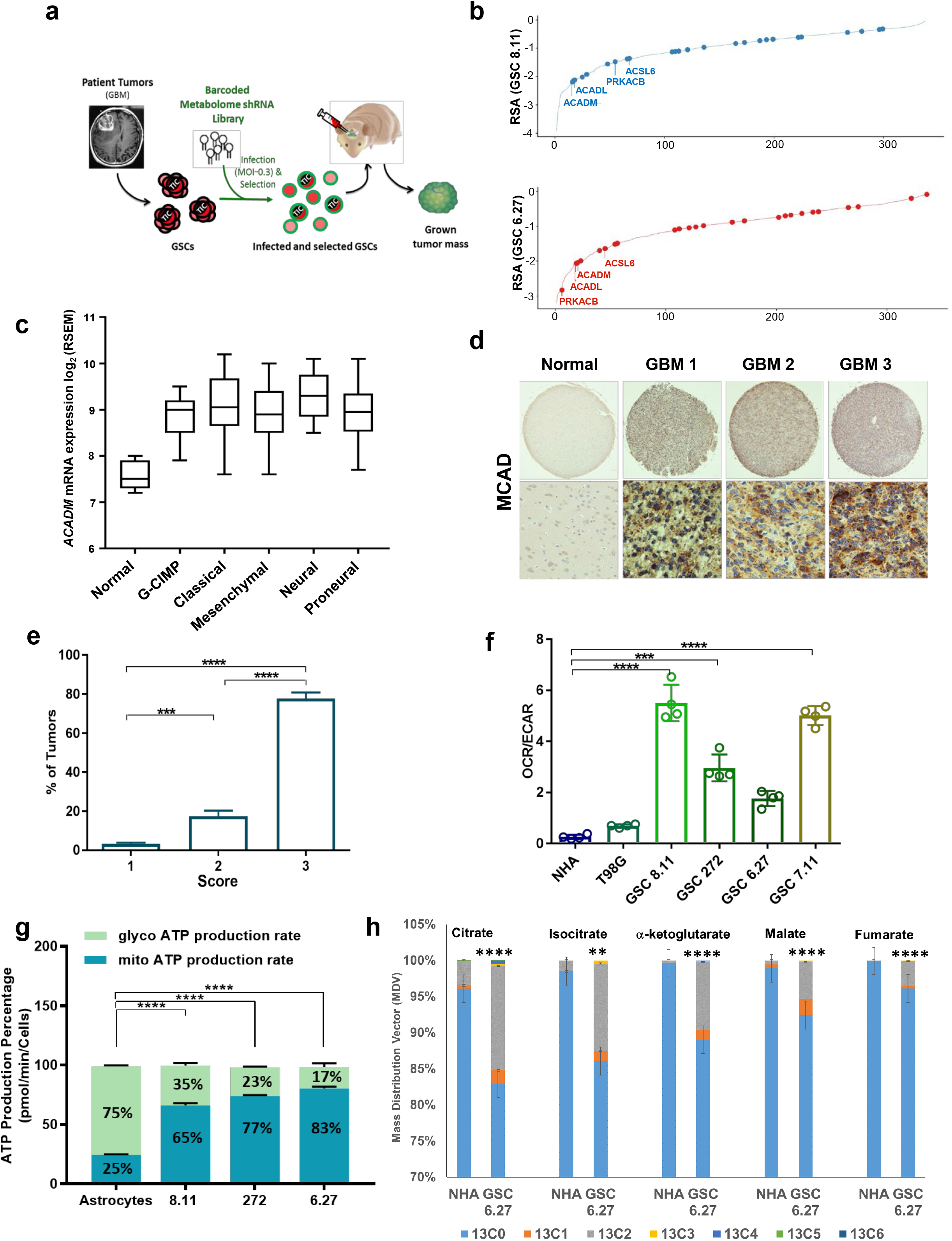
ACADM emerges as a clinically relevant candidate driver of GBM. **a.** Schematics of experimental design for intracranial metabolome shRNA screens in patient-derived glioblastoma stem cells (GSCs). The lentiviral library was transduced at a low multiplicity-of-infection (MOI) (less than one integrant/cell) **b.** Gene-rank analysis highlighting the behavior of genes involved in fatty acid metabolism (*ACADM, ACADL, PRKACB* and *ACSL6*) in *in vivo* screens executed in two independent GSC models: GSC 8.11 and GSC 6.27 (RSA = redundant shRNA activity, logP). **c.** *ACADM* mRNA levels in glioma subtypes vs normal brain (TCGA dataset). **d.** IHC for MCAD on TMA derived from normal brain and GBM tissue. **e.** GBM percentage distribution based on MCAD expression levels in 3 independent TMAs. The scores 1-3 were independently determined using the following scoring system to approximate the percentage of cells positive for staining with the MCAD antibody: 1=0% to 10%, 2=11% to 50%, 3=51% to 100%. Representative tissue scoring is presented in (**d**). Data represent the analysis of three independent TMAs. *P* values generated using oneway ANOVA. \*\*\**P*= 0.0002, \*\*\*\**P* <0.0001. **f.** Bioenergetic profiling of NHA, T98G and GSC lines using Seahorse technology. Basal oxygen consumption rate (OCR) (pMoles/min) and extracellular acidification rate (ECAR) (mpH/min) were used for calculations. Values represent the mean ± SD of four independent experiments. *P* values were generated via one-way ANOVA test. \*\*\**P*= 0.0003, \*\*\*\**P*= 0.0001 **g.** Quantification of energy production in the indicated cell lines by Seahorse XF Real-Time ATP Rate Assay. MitoATP Production Rate and glycoATP Production Rate were calculated from OCR and ECAR measurements under basal conditions. Values are expressed as mean ± SD; *P* values were generated using one-way ANOVA test. \*\*\*\**P* <0.0001 **h.** Isotopologue patterns for incorporation of ^13^C-labeled oleate into TCA cycle intermediates, as measured by LC-MS in NHAs and GSCs in basal conditions. Cells were cultured with ^13^C oleate for 6 hours prior to sample collection. N = 4 biological replicates, error = +/- SD. \*\*\*\**P* <0.0001, ** *P*= 0.0014.

MCAD deficiency is a human inherited autosomal recessive disorder caused by inactivation of the *ACADM* gene. The presence of *ACADM* mutations is screened perinatally, and affected individuals who follow appropriate dietary recommendations can lead normal lives (Schatz and Ensenauer, 2010; Vishwanath, 2016). The most frequent mutation, K304E (~90%), as well as nearly all mutations identified in patients, whether in the catalytic or other domains, disrupt protein folding and prevent active tetrameric MCAD from stabilizing in the mitochondrial matrix (Schatz and Ensenauer, 2010; Vishwanath, 2016). Further, the role of *ACADM* in the pathogenesis of MCAD deficiency has been confirmed by the generation of an *Acadm*-deficient mouse model that shows similar clinical manifestations and histopathological characteristics (Tolwani et al., 2005).

To explore a possible role for *ACADM* in GBM, we queried The Cancer Genome Atlas (TCGA) mRNA dataset and found that *ACADM* is highly expressed across GBM subtypes compared to normal brain (Figure 1C). Through immunohistochemistry analysis, we also determined that MCAD protein levels are elevated compared to normal brain tissue in a GBM tumor microarray series (Figure 1D,E). We selected human astrocytes (NHAs) as a normal tissue control based on previous reports of high *ACADM* mRNA levels relative to other brain-derived cells (Zhang et al., 2014) (Figure S1G), and also given recent evidence that astrocyte-like neural stem cells might be the GBM cell of origin (Lee et al., 2018). We also confirmed lower MCAD levels in NHAs compared to GSCs and the GBM cell line T98G (Figure S2A). A similar trend was observed in the relative expression levels of other enzymes involved in *β*-oxidation (Figure S2B). Consistent with expression levels and previous reports (Vlashi et al., 2011), analysis of mitochondrial bioenergetics demonstrated elevated oxidative metabolism in T98G cells and GSCs compared to NHAs (Figure 1F). Further, we found that GSCs produce ATP predominantly through mitochondrial metabolism (65 to 83%), whereas NHAs generate ATP primarily via glycolysis (~75%) (Figure 1G), thus emphasizing the importance of mitochondrial function for energy generation in GBM. To functionally validate our findings, we performed metabolic flux analysis (MFA) using ^13^C-oleate as the substrate. Strikingly, these data demonstrated a significant enrichment of ^13^C in TCA cycle metabolites in GSCs compared to NHAs (Figure 1H, Figure S2C), thus supporting that oxidative metabolism and FAO are both enhanced in GBM.

To further investigate the relevance of MCAD in this context, we validated two independent *ACADM-* targeting RNA interference (shRNA) constructs for their ability to down-regulate MCAD protein expression (Extended Data Fig. 3A). We found that MCAD depletion dramatically impaired anchorage-independent growth in multiple GSCs representing different GBM subtypes (Figure 2A-B, Figure S3B). Similar effects on GSC growth were obtained by *ACADM* deletion through CRISPR-Cas9 editing using three independent sgRNA constructs (Figure S3C-E). To better understand the effects of *ACADM* down-regulation on GSCs, we evaluated apoptosis in a time course experiment, which demonstrated that the growth arrest induced by *ACADM* ablation precedes a significant increase in apoptotic cell death (annexin V positivity) detected approximately 7 days after the end of selection in all GSCs tested (Figure 2C; Figure S4A,B). Indeed, no significant effect on cell viability was observed at earlier time points (72 hours) (Figure 2C, Figure S4A). Similar effects on cell growth and sphere-forming efficiency were observed upon treating GSCs with spiropentaneacetic acid (SPA), a compound known to specifically inhibit MCAD activity (Tserng et al., 1991) (Figure 2D; Figure S4C,D).

**Figure 2.**
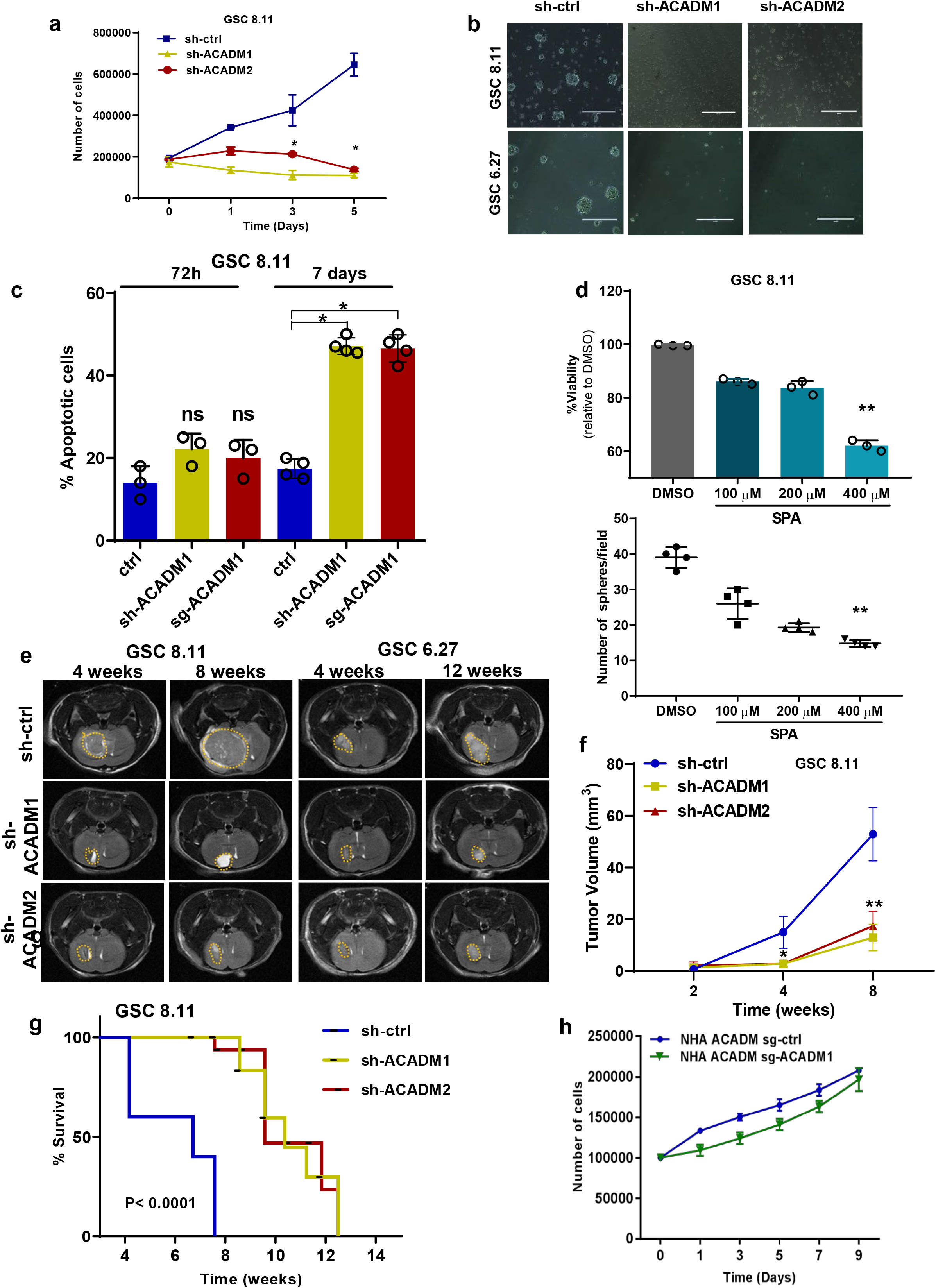
MCAD is essential for GSC proliferation and tumorigenesis. **a.** Five-day growth curve of GSC 8.11 cells upon shRNA *ACADM* silencing. Day 0 was defined as 48 hours post-puromycin selection. Values represent the mean ± SD of three independent experiments. *P* values were generated using one-way ANOVA. \**P* ≤0.02. **b.** Representative optical microscopy images of GSC tumor spheres 4 days after puromycin selection. Scale bar = 100 μm. **c.** Quantification of apoptosis in GSC 8.11 infected with anti-*ACADM* or scrambled shRNA by Annexin V-FITC/PI. Staining was evaluated by flow cytometry at 72 hours and on day 7 after puromycin selection. Values represent the mean ± SD of three or four independent experiments; *P* values were generated using one-way ANOVA test. \**P* < 0.04. **d.** (Top) Cell viability assessed by Trypan Blue exclusion of GSC 8.11 treated with indicated concentration of SPA for 96 hours. Data represent mean ± SEM of three biologically independent replicates. *P* values were generated using one-way ANOVA. \*\**P* = 0.0065. (Bottom) Dot plot showing GSCs 8.11 sphere formation efficiency (SFE) after 48-hour exposure to SPA at indicated concentrations. DMSO was used as control. Values represent the mean ± SD of four independent experiments; *P* values were generated using one-way ANOVA test. \*\**P* = 0.0011. **e.** MRI images of tumor progression after implantation of GSCs 8.11 or 6.27 at 4 and 8 weeks for GSCs 8.11 and at 4 and 12 weeks for GSC 6.27. **f.** Quantification of tumor progression after implantation of GSC 8.11 as measured by MR volumetry (n=8 mice per group). *P* values were generated using one-way ANOVA. \**P*= 0.04, \*\**P* = 0.0082. **g.** Kaplan-Meier survival analysis after implantation of GSCs 8.11. For sh-ctrl, sh-*ACADM1*, sh-*ACADM2*, n=8 mice per group. *P* values were generated using log-rank test. **** *P* < 0.0001. **i.** Growth curve of normal human astrocytes (NHA) infected with sgRNA targeting either *ACADM* or GFP. Values represent the mean ± SD of three independent experiments. **h.** Growth curve of NHAs infected with sgRNA targeting either *ACADM* or GFP. Values represent the mean ± SD of three independent experiments.

To determine the effect of MCAD loss *in vivo*, GSC 8.11 and GSC 6.27 cells harboring *ACADM-* or nontargeting shRNA constructs were implanted into the mouse forebrain, and tumor growth was monitored by luciferase and magnetic resonance imaging (MRI). MCAD depletion dramatically attenuated tumor growth and significantly extended survival time (Figure 2E-G; Figure S5A-C). Similarly, MCAD depletion in tumors established through intracranial injection of GSC 8.11 cells carrying doxycycline-inducible shRNA constructs resulted in significant retardation of tumor growth (Figure S5D). We next generated *ACADM-* deleted NHAs using our validated sgRNA guides. Interestingly, neither the cytotoxic nor the antiproliferative effects observed in GSCs were observed in NHAs (Figure 2H; Figure S5E), suggesting that MCAD dependency may be a metabolic vulnerability unique to malignant cells.

Given that GSC energetics seem to be heavily reliant on oxidative metabolism (Figure 1F,G), we characterized the early effects of MCAD downregulation on mitochondria between 48 and 72 hours post-MCAD depletion (shRNA/sgRNA ablation). First, we acquired transmission electron microscopy (TEM) images revealing that MCAD depletion resulted in mitochondria with darker matrices and swollen cristae surrounded by vacuolar structures (Figure 3A; Figure S6A). Next, a bioluminescence assay uncovered an overall decrease in ATP content in MCAD-depleted GSCs versus controls (Figure 3B). Analysis of oxygen consumption rate (OCR) in GSCs and NHAs revealed significant decreases in basal respiration and reserve respiratory capacity in *ACA33DM*-deleted GSCs (Figure 3C; Figure S6B), whereas *ACADM* deletion did not affect OCR in NHAs (Figure 3D). The depletion of the acetyl-CoA pool and TCA intermediates upon *ACADM* down-regulation (Figure 3E,F; Figure S6C), together with an increased contribution of carbon skeletons derived from fatty acids to central carbon metabolism in GBM (Figure 1H), confirm the critical dependence of GBM cells on oxidative metabolism, largely fueled by FAO, and the role of MCAD to support mitochondrial function in this context.

**Figure 3.**
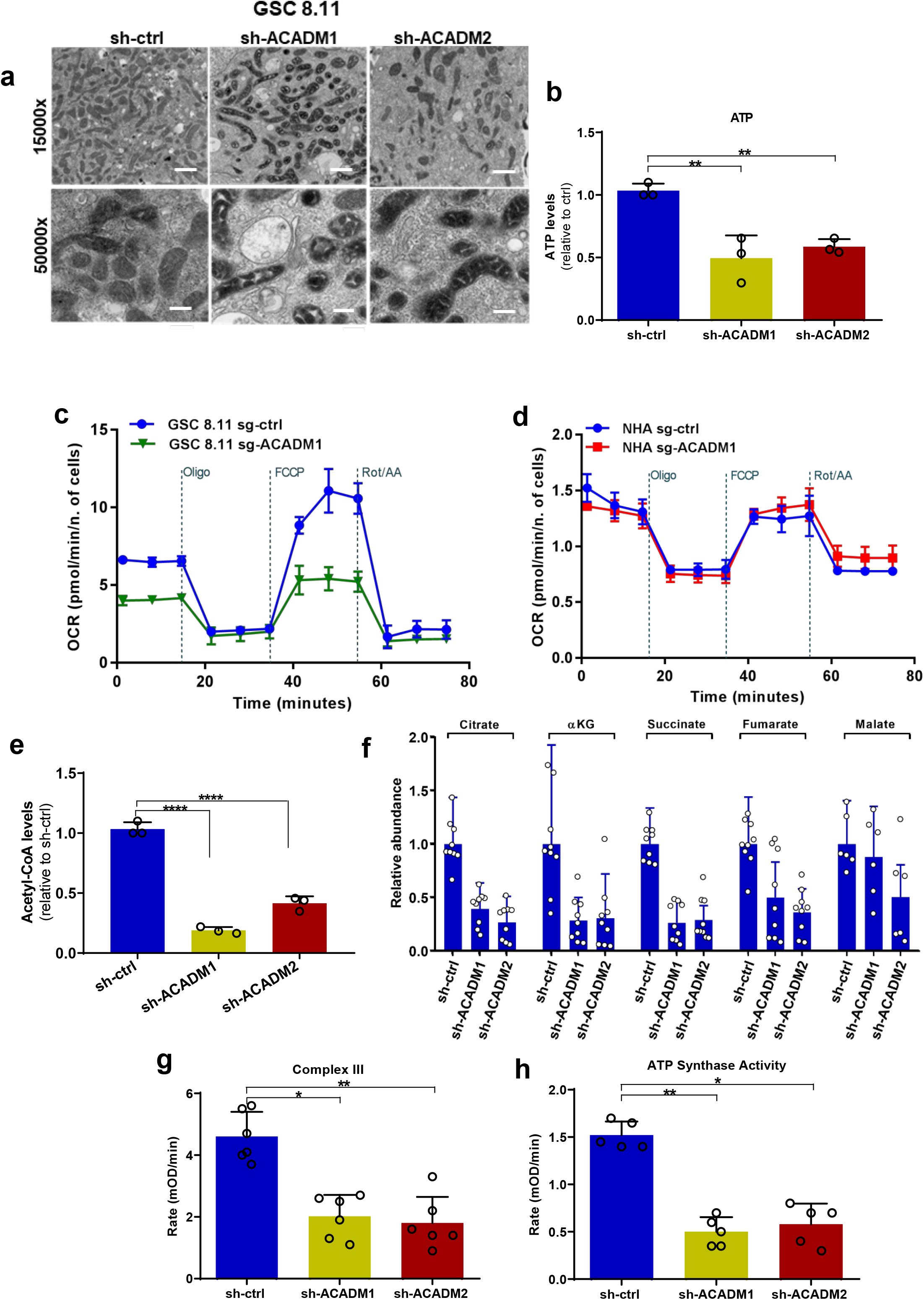
ACADM silencing causes mitochondrial failure in GSCs. **a.** Transmission electron microscopy images of mitochondria in GSC 8.11 upon MCAD silencing (magnification x15,000 and 50,000; scale bar= 1 μm, 300 nm) **b.** Measurements of mitochondrial ATP levels using a bioluminescence assay. Values are expressed as mean ± SD of three independent experiments; *P* values were generated using one-way ANOVA test. \*\**P* < 0.006. **c., d.** Oxygen consumption rate (OCR) measured in GSC 8.11 (**c**) and in normal human astrocytes (NHA) (d) following MCAD silencing by metabolic flux assay before and after the addition of oligomycin, FCCP and rotenone/antimycin to derive several parameters of mitochondrial respiration. Values represent the mean±SD of one representative experiment with n = 4 technical replicates. Experiments were repeated three times with similar results (see also ED Fig. 6b). **e.** Intracellular acetyl-CoA levels in GSC 8.11 after *ACADM* silencing as assessed by fluorometric assay. Values are expressed as mean ± SD of three independent experiments; *P* values were generated using one-way ANOVA test. \*\*\*\**P* = 0.0001. **f.** Relative abundance of metabolites in glycolysis in GSC 8.11, as detected by LC-MS analysis. Values are expressed as mean ± SD of three independent experiments. **g.** Mitochondrial electron transport chain complex III activity in GSC 8.11 as measured by colorimetric assay. Values are expressed as mean ± SD of six independent experiments. *P* values were generated using one-way ANOVA test. \**P* = 0.01, \*\**P* = 0.0079. **h.** Mitochondrial complex V (ATP synthase) activity in GSC 8.11 as measured by colorimetric assay. Values are expressed as mean ± SD of five independent experiments; *P* values were generated using one-way ANOVA test \**P* = 0.0313, \*\**P* = 0.0071.

To distinguish whether the observed toxicity of MCAD inactivation is triggered by mitochondrial dysfunction or by the energy deprivation resulting from the inability of GSCs to oxidize fatty acids, we grew *ACADM*-silenced GSCs in medium supplemented with acetate, which is a source of carbon molecules that can bypass MCAD activity by directly fueling the TCA cycle (acetyl-CoA), or with long-chain fatty acids (LCFAs; e.g. linoleic acid) that are not metabolized by MCAD. Neither acetate nor LCFA supplementation rescued proliferation in MCAD-depleted GSCs (Figure S6D,E), which strongly suggests that MCAD ablation, in addition to depleting energy substrates, may compromise overall mitochondrial function. In support of this hypothesis, we observed significant impairment of mitochondrial Complex III and Complex V (ATP synthase) activity in MCAD-depleted versus-proficient cells (Figure 3G,H).

The physiological role of MCAD is to degrade MCFAs that freely diffuse into cells (Marten et al., 2006). Thus, we hypothesized that decreased MCAD function may result in a toxic accumulation of lipids that are normally diverted to energy production, triggering mitochondrial failure. Consistent with this hypothesis, quantification of lipids in GSCs using Oil Red O lipid staining revealed massive accumulation of lipid droplets *in vitro* as early as 48 hours after puromycin selection in *ACADM*-silenced cells, as well as in GSCs pharmacologically treated with SPA (Figure 4A; Figure S7A). Similarly, MCAD deficiency increased overall free fatty acid content, as assessed by a colorimetric assay (Figure 4B). Lipid accumulation was also confirmed *in vivo* in tumor remnants after 30 days of doxycycline-induced *ACADM* silencing (Figure 4C; Figure S5D). To test whether the accumulation of fatty acids is causally correlated with the MCAD-deficient phenotype, we ablated MCAD in T98G cells kept in medium supplemented with normal or lipoprotein-deficient serum (LPDS). In these lipid-depleted conditions, MCAD ablation did not result in accumulation of lipid droplets. Moreover, cell proliferation, as well as viability, albeit reduced, did not appear significantly impacted (Figure 4D,E; Figure S7B).

**Figure 4.**
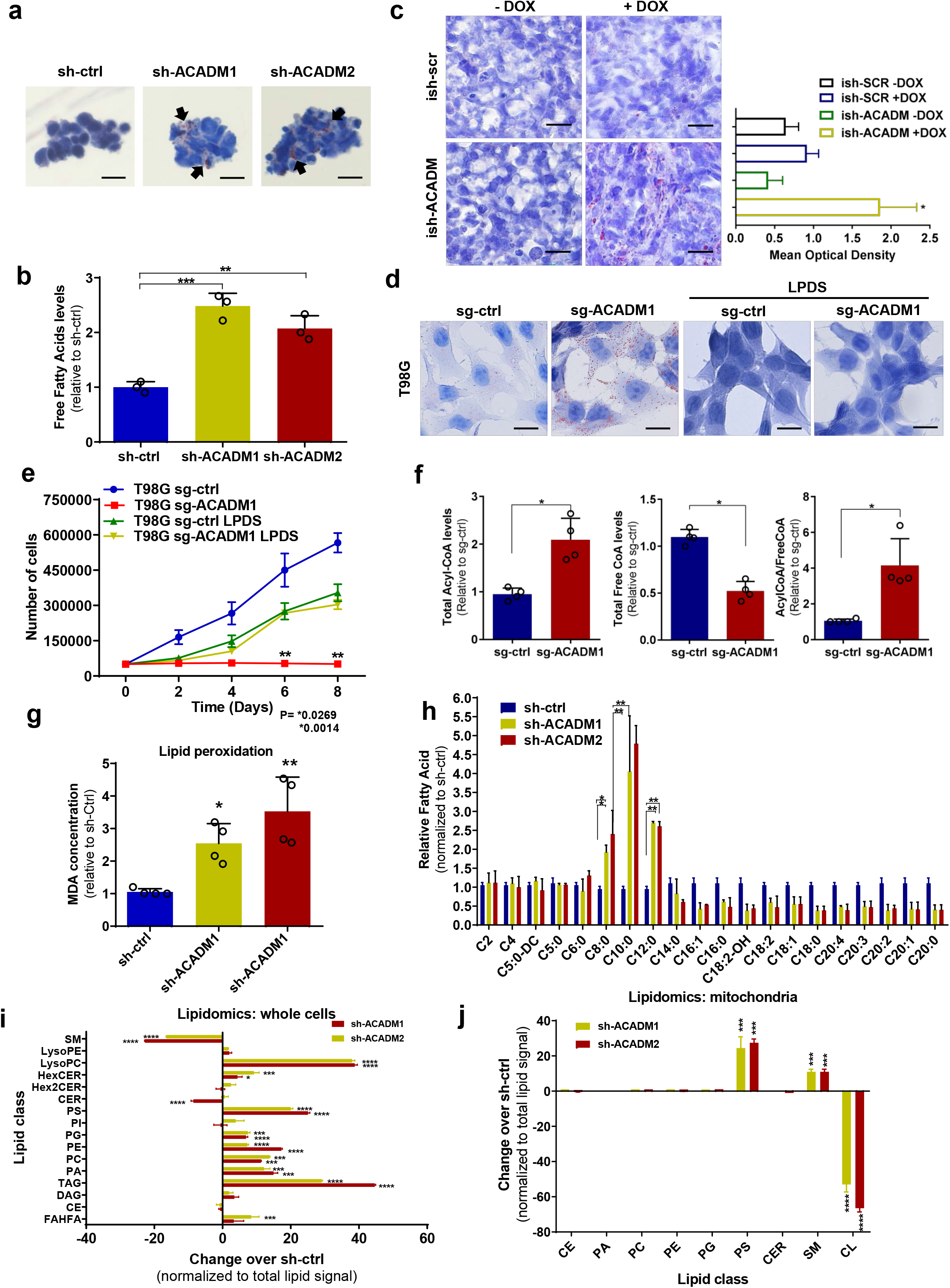
Accumulation of MCFAs induces toxic alterations of lipid metabolism in GSCs. **a**. Oil Red O staining in GSC 8.11 cells infected with anti-*ACADM* or non-targeting shRNA *in vitro*. Black arrows indicate sites of lipid accumulation. Scale bar = 20 μm **b.** Colorimetric determination of free fatty acids from GSC whole-cell extracts. Values represent the mean ± SD of three independent experiments. *P* values were generated using one-way ANOVA test. \*\**P* = 0.0061; \*\*\**P* = 0.0003. **c.** Oil Red O staining in xenograft tumor tissues derived from GSC 8.11 infected with inducible shRNA constructs. Doxycycline was administered approximately 20 days after cell implantation. Oil Red O staining quantification of tumor tissues shown in (**c**) was obtained using ImageJ software analysis. *P* values were generated using one-way ANOVA. \**P* = 0.033. **d., e.** Oil Red O staining (**d**) and growth curve (**e**) of *ACADM* wild type or null T98G cells grown in medium containing FBS or LPDS. Cells were selected with puromycin for 48 h prior to starting the experiment. Oil Red O staining in cells after 48-h puromycin selection. Scale bar in d = 10 μm. Data in (**e**) represent the mean ± SD of three biologically independent replicates. *P* values were generated using one-way ANOVA. \*\**P* ≤ 0.007. **f.** Colorimetric analysis of acyl-CoA species in MCAD-deficient GSC 8.11 cells. Data represent the mean ± SEM of four biologically independent replicates. *P* values were generated using one-way ANOVA. \**P* <0.03. **g.** Quantification of lipid peroxidation determined by measuring the production of malondialdehyde (MDA) using the Colorimetric Microplate Assay for Lipid Peroxidation Kit. Data represent the mean ± SD of four biologically independent replicates. *P* values were generated using one-way ANOVA. \**P* = 0.02, \*\**P* = 0.001. **h.** Quantitative LC-MS/MS lipid profiling of FFA content of *ACADM* wild type or null GSC 8.11 cells. Data represent mean ± SEM of three biologically independent replicates. *P* values were generated using one-way ANOVA. \**P* < 0.03; \*\**P* <0.004. **i., j.** Relative amount of total lipid classes measured by mass spectrometry of whole-cell extracts (**i**) or mitochondria (**j**) from GSC 8.11 cells infected with shRNA targeting *ACADM*. Data are reported as fold change over control cells infected with non-targeting shRNA. Mean values ± SD of three biologically independent replicates. *P* values were generated using one-way ANOVA. \*\*\**P* = 0.001; \*\*\*\**P* = 0.0001. SM, sphingomyelin; LPE, lyso-phosphatidylethanolamine; LPC, lyso-phosphatidylcholine; HexCER, hexosylceramide; Hex2CER, dihexosylceramide; CER, ceramide; PS, phosphatidylserine; PI, phosphatidylinositol; PG, phosphatidylglycerol; PE, phosphatidylethanolamine; PC, phosphatidylcholine; PA, phosphatidic acid; TAG, triacylglycerol; DAG, diacylglycerol; CE, cholesteryl ester; FAHFA, fatty acid ester of hydroxyl fatty acids.

Our data strongly suggest that lipid accumulation contributes directly to the proliferation defect observed in MCAD-deficient GSCs. To understand mechanistically how lipid accumulation may exert toxicity, we investigated the effects of MCAD depletion on coenzyme A (CoA) pools in GSCs, as conjugation of fatty acids to CoA as acyl-CoA is required for fatty acids to migrate into mitochondria and undergo oxidation. Compared with wild type cells, MCAD-depleted cells had increased acyl-CoA and decreased free-CoA pools (Figure 4F). Further, peroxidation of accumulated lipids was increased in the absence of MCAD (Figure 4G), which would also be expected to impact cellular structures and functions. This important finding suggests that activated fatty acids may enter and accumulate in the mitochondria.

To characterize the type and size of lipids accumulated in MCAD-depleted cells, we compared free fatty acid profiles of *ACADM*-silenced versus control GSCs by gas chromatography mass spectrometry (GC-MS). Here, we observed an accumulation of MCFAs and a decrease in LCFAs (Figure 4H), consistent with a block in the metabolic pathway at the level of MCFA degradation. Next, we profiled acyl-carnitines by liquid chromatography mass spectrometry (LC-MS) in GSCs grown in medium supplemented with unmodified or uniformly carbon-13 (U^13^C18)-labeled oleate. This experiment revealed an increase in medium-chain acyl carnitine species that would be expected to accumulate in the mitochondria (Figure S7C). Thus, similar to observations in patients affected by MCAD deficiency (Smith et al., 2010), the accumulation of MCFAs is observed both as free fatty acid and acyl-carnitine species, suggesting that GSCs, to some extent, continue to engage in FAO (e.g., oleate), even upon attenuated MCAD activity.

MCFAs have been shown to induce apoptosis in some cancer models (Fauser et al., 2013; Lappano et al., 2017) and, based on the dramatic accumulation of MCFAs in GSCs upon MCAD depletion, we speculated that MCFAs may be directly causing toxicity. Indeed, acute treatment of GSCs with varied-length fatty acid species (between 4 and 16 carbons) revealed that C10 and C12 MCFAs negatively impacted cell viability, whereas treatment with the shorter or longer fatty acids tested did not change viability (Figure S7D). Interestingly, acute MCFA exposure induced an increase in ROS levels that was partially rescued when cells were concomitantly treated with antioxidants such as GSH-EE (Figure S7E,F). Because lipid peroxidation products have been associated with ferroptosis (Yang and Stockwell, 2016), an iron-dependent form of programmed cell death, we silenced *ACADM* in GSCs in the presence of ferrostatin-1, an iron chelating agent. Propidium iodide staining did not reveal any improvement in viability in *ACADM*-silenced GSCs upon ferrostatin-1 treatment (Figure S7G), which suggests that the toxicity exerted by fatty acid accumulation is iron independent. Taken together, our data support a model wherein MCAD inhibition negatively affects cell viability through the accumulation of toxic species as a consequence of impaired fatty acid degradation.

Next, to investigate the effects of MCAD depletion on cellular lipids more broadly, we conducted LC-MS-based analysis of lipid classes on whole-cell extracts and on mitochondria isolated from GSCs. Triacylglycerols (TAG) and some phospholipid (PL) classes, such as lysophosphatidylcholine (LysoPC), were significantly increased in whole-cell extracts from MCAD-depleted cells compared with MCAD-proficient controls (Fig. 4I). This indicates that profound perturbations in the cellular content of complex lipids may reflect an adaptation by which MCAD-inhibited GSCs attempt to accommodate the accumulating MCFA species. Indeed, phosphatidylserine and sphingomyelin levels were approximately 20- and 10-fold higher, respectively, in MCAD-depleted versus control GSCs analyzed by LC-MS lipidomics (Figure 4J). The most dramatic change detected in mitochondria was in cardiolipin, which decreased over 50-fold in MCAD-deficient cells compared to controls (Figure 4J). Cardiolipin is located in the inner mitochondrial membrane, where it plays a crucial role in mitochondrial bioenergetics and regulates the efficiency of the electron transport chain (ETC). Interestingly, it is well established that cardiolipin is the phospholipid most susceptible to oxidative stress due to its composition, which is enriched in unsaturated fatty acids (Paradies et al., 2014).

To evaluate whether these changes in mitochondrial lipids may relate to mitochondrial dysfunction, we measured ROS levels in MCAD-deficient GSCs. MCAD depletion by shRNA or SPA treatment altered the cells’ redox state, as evidenced by increased ROS levels as well as decreased ROS scavengers and NADPH levels in all GSCs tested (Figure 5A-D; Figure S8A-C). This correlates with our previous finding of high levels of lipid peroxidation in this context (Figure 4G). We also found high levels of 8-oxoguanine and 4-HNE-lysine, two indicators of oxidative damage and lipid peroxidation, which supports that MCAD depletion similarly affects redox balance in tumors *in vivo* (Figure 5E; Figure S8D). To investigate the impact of redox state on viability, we grew *ACADM*-silenced GSCs in suspension culture in the presence of a cell-permeable form of glutathione (GSH-ethyl ester, GSH-EE). GSH-EE exposure transiently rescued the growth inhibition observed upon MCAD depletion; however, it did not enable MCAD-depleted spheres to expand to the same extent as control spheres at later time points (Figure 5F,G; Figure S8E,F). Thus, despite the central role of ROS in driving the observed phenotype, our data convincingly demonstrate that oxidative damage is only one aspect of how deficient MCAD function affects GSCs. In strong support of this, culturing MCAD-depleted GSCs in LPDS medium preserved the proliferative phenotype (Figure 4E) and ablated the increase in ROS induced by *ACADM* silencing (Figure S8G). Taken together, our functional characterization of MCAD in glioblastoma models uncovers a novel, protective role wherein MCAD function is required for continuous degradation of molecular species that would otherwise accumulate and cause mitochondrial dysfunction, oxidative stress and, eventually, cell death.

**Figure 5.**
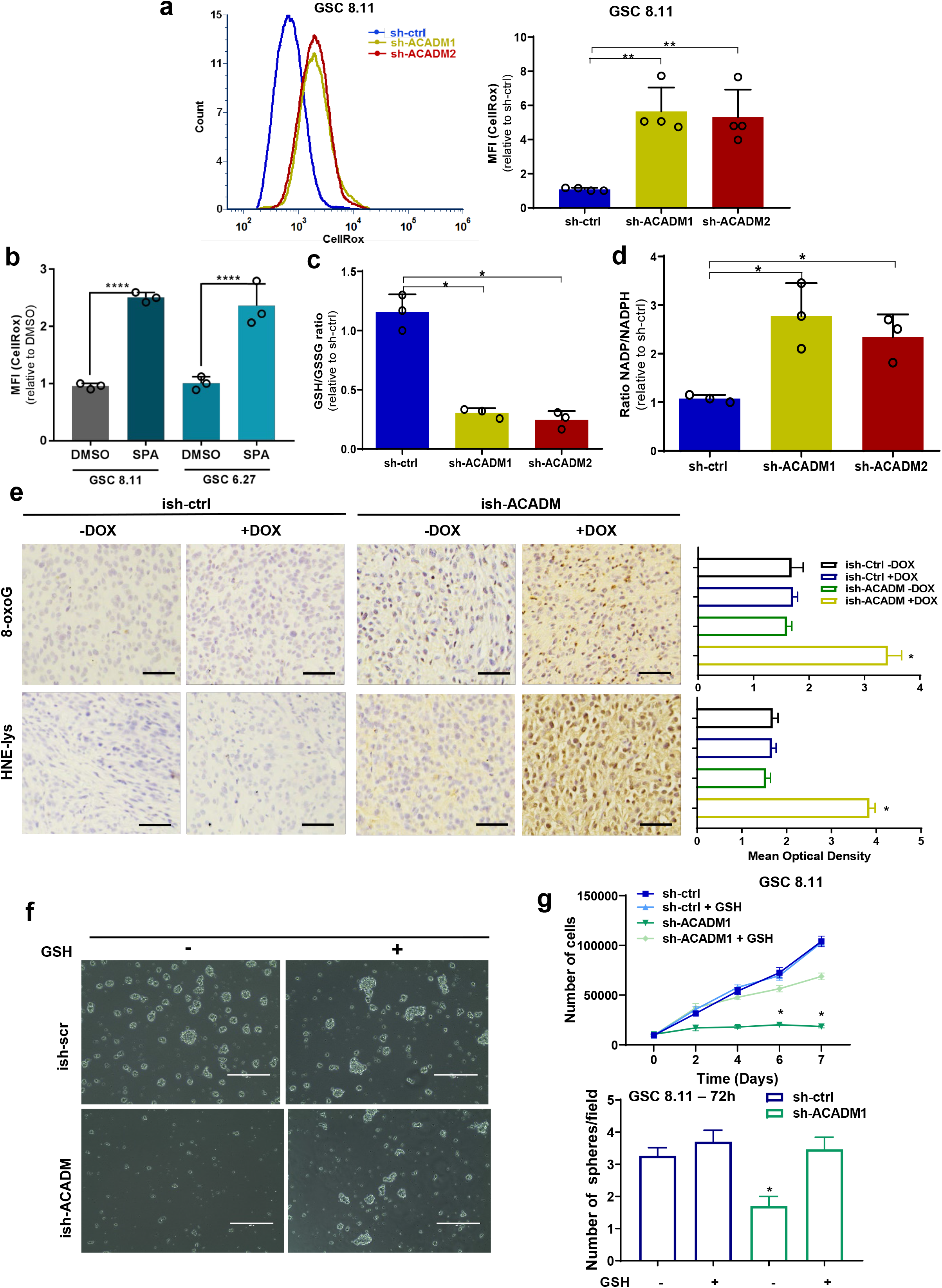
MCAD knockdown triggers ROS-related damage in GSCs in vitro and in vivo. **a.** ROS production as measured by CellROX Green in flow cytometry (left) and quantification of fluorescence intensity (right) in GSC 8.11 harboring anti-*ACADM* or non-targeting shRNA. Values represent the mean ± SD of four independent experiments. *P* values were generated using one-way ANOVA test. \*\**P* < 0.002. **b.** ROS production quantification by CellRox Green staining intensity in GSC 8.11 and 6.27 upon 72-hour exposure to SPA (400 μM) or DMSO. Values represent the mean ± SD of three independent experiments. *P* values were generated using one-way ANOVA test. \*\*\*\**P*= 0.0001. **c. d.** GSH/GSSG and NADP/NADPH ratios in GSC 8.11 measured by colorimetric assay. **(c)** \**P* ≤ 0.03; **(d)** \**P* ≤ 0.03. Values represent the mean ± SD of three independent experiments. *P* values were generated using one-way ANOVA test. **e.** Immunostaining for 8-oxoguanine (showing oxidized DNA, top panels) or HNE adducts (showing lipid peroxidation and oxidative protein damage, bottom panels) and relative quantification of GSC 8.11-derived tumors from experiment shown in Extended Data Fig. 5D; scale bar 50 μm. Quantification was conducted with ImageJ software, six images per condition, where each condition was represented by three biological replicates. Values are expressed as mean ± SD; *P* values were generated using one-way ANOVA test. \**P* ≤ 0.02. **f.** Representative images of the rescue effect of GSH-ethyl ester (GSH-EE) on GSC 8.11 tumor spheres with doxycycline-induced MCAD knockdown (scale bar 400 μm). **g.** (Top) Growth curve of GSC 8.11 cells infected with *ACADM* or non-targeting shRNA grown in the presence or absence of GSH-EE. Values are expressed as mean ± SD; *P* values were generated using one-way ANOVA test. \**P* ≤ 0.03; (bottom) Quantification of the number of spheres formed by GSC 8.11 and GSC 6.27 harboring *ACADM* or nontargeting shRNAs grown for 72 hours or 7 days in the presence or absence of GSH-EE, as indicated. Values represent the number of spheres per field expressed as mean ± SD. *P* values were generated using oneway ANOVA test. \**P*= 0.04.

In conclusion, here we describe a previously unappreciated metabolic dependency in glioblastoma that was identified through a functional genomics screen conducted in orthotopic, GSC-derived models of GBM. Previous reports have indicated that the upregulation of FAO confers a survival advantage to GBM cells, at least in part by fueling oxidative metabolism to sustain cellular energetic requirements (Lin et al., 2016). The significance of FAO to fuel cancer cell survival beyond brain cancer is rapidly gaining recognition (Carracedo et al., 2013). Several studies have reported that blocking LCFA oxidation, primarily through CPT1 inhibition, exerts robust anti-tumor effects in pre-clinical models, but CPT1-targeted therapies have not yet proven to be clinically viable due to significant toxicity (Aires et al., 2010; Cabrero et al., 2003; Haynie et al., 2014; He et al., 2012; Sayed-Ahmed et al., 2000). This work expands our understanding of the dependence of glioblastoma cells on FAO beyond supporting cancer growth energetic requirements and demonstrates that the sustained influx of fatty acids in glioblastoma cells can become toxic when lipids are not properly metabolized, identifying a novel protective role for MCAD in GBM. In fact, our findings demonstrate that accumulated MCFAs are particularly damaging to GSCs compared to fatty acids of different chain lengths, supporting the strategy that blocking FAO at the level of MCFA oxidation may be an effective approach to induce a catastrophic cell death in GBM. Indeed, targeting MCAD may have advantages over inhibitors of CPT1, because partial oxidation of LCFA to MCFA may contribute to the overall accumulation of toxic lipid species in contexts where MCFA metabolism is inefficient. Given the toxicity associated with C10 and C12 accumulation and the lack of a feedback mechanism to prevent continued MCFA accumulation, we posit that, in a lipid-rich environment such as the brain, overexpression of MCAD or enzymes capable of diverting potentially harmful lipid pools to energy production may confer a significant proliferation advantage to cancer cells and explain the high reliance of GSCs on sustained fatty acid metabolism. Therefore, targeting MCAD in fatty acid-embedded tumor types may represent a unique and therapeutically exploitable vulnerability. This, along with the well-documented ability to manage congenital MCAD deficiency syndrome through dietary adjustment, suggests that there may be a viable therapeutic window to develop an MCAD-targeted therapy in patients affected by glioblastoma and potentially other conditions.

## METHODS

### Human samples, primary cells and cell lines

Glioblastoma stem cells (GSCs) were isolated from specimens from patients with glioblastoma, who had undergone surgery at the University of Texas MD Anderson Cancer Center (UTMDACC). The diagnosis of glioblastoma was established by histological examination by the WHO (World Health Organization) classification. Samples derived from patients were obtained with the consent of patients, under an ethically approved Institutional Review Board protocol LAB04-0001 chaired by F. F. Lang (UTMDACC). Tumor specimens were dissociated and resuspended in Dulbecco’s modified Eagle’s medium/F12 (DMEM/F12, Gibco) supplemented with B27 (×1, Invitrogen), bFGF (20 ng ml-1 Peprotech), and EGF (20 ng ml-1, Peprotech). Cells were cultured as neurospheres and passaged every 5–7 days on the basis of sphere size. Human astrocytes were purchased from Sciencell (#1800) and grown in Astrocyte Medium (#1801) according to instructions. T98G cells were obtained through ATCC and grown under standard tissue-culture conditions.

### GSCs treatments, proliferation and sphere formation assays

Sodium butyrate (156-54-7), sodium hexanoate (C4026), sodium octanoate (C5038), sodium decanoate (C4151), and sodium dodecanoate (L9755) were obtained from Sigma-Aldrich. BSA-Palmitate (102720-100) was purchased from Agilent Technologies. Spiropentaneacetic acid (SPA) was synthesized by the Institute of Applied Cancer Science Chemistry Core Facility at MD Anderson as previously described^33^. For each treatment, GSCs were disaggregated and 50000 cells were plated in 6-well plates. Cells were disaggregated at different time points (as indicated in Figures); cell number and viability were determined by Trypan Blue exclusion assay. For sphere formation assays, approximately 2 x 10^4^ cells from disaggregated GSC spheres were resuspended in a 0.8% methylcellulose semisolid DMEM/F12 and seeded in 6-well plates. At different time points, the spheres were counted in at least 4 fields per well.

### Library Design and Construction

A custom library was composed by 338 genes specifically belonging to key metabolic pathways (KEGG, see Figure S2D). This library was constructed by using chip-based oligonucleotide synthesis and cloned into the pRSI16 lentiviral vector (Cellecta) as a pool. The shRNA library is constituted by 338 genes with a coverage of 10 shRNAs/gene. The shRNA includes two G/U mismatches in the passenger strand, a 7-nt loop and a 21-nt targeting sequence. Targeting sequences were designed using a proprietary algorithm (Cellecta). The oligo corresponding to each shRNA was synthesized with a unique molecular barcode (18 nucleotides) for measuring representation by next generation sequencing.

### *In Vivo* shRNA Screens

*In vivo* shRNA screens were performed using adapted procedures previously described in (Carugo et al., 2016). In brief, concentrated lentiviral particles (TU, transducing units) from libraries or single plasmids were either purchased by Cellecta or produced by transfecting 293T cells according to the protocol in the Cellecta User Manual. Precisely calculated lentiviral particles, together with 2 μg/mL polybrene (Millipore), were added to single-cell dissociated GSCs to achieve a multiplicity of infection (MOI) = 0.3. Forty-eight hours after infection, the medium was replaced and puromycin (2 μg/mL) was added for 96 hours. For *in vitro/in vivo* validation studies, GSCs were infected at MOI = 3, with single shRNA knocking down specific target genes. Transduction efficiency was determined sample by sample as the percentage of GFP positive cells 2 days after infection as measured by FACS analysis. For the *in vivo* experiments, each intracranial injection was performed with 1 × 10^6^ cells to ensure a coverage of ~300 cells/barcode. Upon brain collection, tumors displaying green fluorescence signal were precisely dissected, weighted and quickly snap frozen. Genomic DNA extraction, barcode amplification and next-generation sequencing were performed accordingly to Cellecta User Manual for shRNA libraries processing (details about PCR primers, PCR conditions and reads counting in (Carugo et al., 2016).

### Bioinformatics analysis - Hit identification

Read counts were normalized for the amount of sequencing reads retrieved for each sample by using library size normalization (to 1 million reads). For each sample (log2) fold-change, was calculated compared to the reference pellet before injection. A summary measure per condition was derived using median of quantile transformed log2 FC across replicates. Thereafter, a modified version of the RSA algorithm was used to derive the gene level summary measure per condition. Specifically, we controlled for the factor that at least 2 hairpins were used when calculating the minimum p-value (in RSA). In addition, hairpins which ranked above luciferase controls were not used in choosing the minimum p-value. Quantile rank of luciferase controls barcodes was evaluated across all experiments. On average, luciferase barcodes ranked >0.6 (on the quantile transformed log2fc scale). Therefore, hairpins with quantile transformed log2fc > 0.6 were not used for gene-level RSA scores (log value) (Birmingham et al., 2009).

### Animal Studies

Male athymic nude mice (*nu/nu*) were purchased from The Jackson Laboratories. All procedures performed in this study were approved by the University of Texas MD Anderson Cancer Center. All animal manipulations were performed in the veterinary facilities in accordance with institutional, state, and federal laws and ethics guidelines under an approved protocol. Intraperitoneal (IP) injections of ketamine (100mg/kg)/xylazine (10mg/kg) were used to anesthetize animals in all experiments. For intracranial xenografts of GSC 8.11 and 6.27, 10^6^ cells expressing a PLX304-mCherry-LUC vector (5 μl cell suspension) were implanted using a guide screw and a multiport microinfusion syringe pump (Harvard Apparatus, Holliston, MA) (Lal et al., 2000; Nakamizo et al., 2005). For subcutaneous experiments, GSCs were injected subcutaneously (100 μl cell suspension) into the left flank. For all bioluminescence imaging, d-luciferin (150 mg kg^-1^) was administered by subcutaneous injection to mice 10 min before imaging. In all experiments, mice were monitored daily for signs of illness and were euthanized when they reached endpoints.

For Magnetic Resonance Imaging (MRI), a 7T Bruker Biospec (BrukerBioSpin), equipped with 35mm inner diameter volume coil and 12 cm inner-diameter gradients, was used. A fast acquisition with relaxation enhancement (RARE) sequence with TR/TE of 2,000/39 ms, matrix size 256×192, Resolution was 156uM, 0.75 mm slice thickness, 0.25 mm slice gap, 40 x 30 cm FOV, 101 kHz bandwidth, 4 NEX was used for acquired in coronal and axial geometries a multi-slice T2-weighted images. To reduce the respiratory motion the axial scan sequences were respiratory gated. The brain lesions’ volumes were quantified manually using Image J software. All animal imaging, preparation and maintenance was carried out in accordance with MD Anderson’s IACUC policies and procedures.

### ETC activity

The Mitochondria Isolation Kit (Abcam ab110170) was used to isolate mitochondria from MCAD silenced and control GSCs, which were then used to test the activity of Complex III and V. Activity of mitochondrial complex III was analyzed using the Mitochondrial Complex III Activity Assay Kit (Biovision K520-100). Cytochrome C was added to the reaction and its reduction through the activity of Complex III was measured at 550 nm. ATP synthase activity was measured by using the ATP Synthase Specific Activity Microplate Assay Kit (Abcam ab109716). The antibody for ATP Synthase was precoated in the wells of the microplate, and samples containing 50 μg of mitochondrial extracts were added to wells. In this assay, the conversion of ATP to ADP by ATP synthase was coupled to the oxidative reaction of NADH to NAD+. The formation of NAD+ resulted in a decrease in absorbance at 340 nm. Subsequently, in these same wells, the quantity of ATP synthase was determined by adding an ATP synthase specific antibody conjugated with alkaline phosphatase. This phosphatase changed the substrate pNPP from colorless to yellow (OD 405 nm) in a time dependent manner proportional to the amount of protein captured in the wells. Absorbance was read by a Benchmark microplate reader (Bio-Rad). All tests were done in triplicate.

### ATP quantification

ATP was quantified by using the luminescent ATP detection assay from Abcam (ab113849). The assay quantifies the amount of light emitted by luciferin when oxidized by luciferase in the presence of ATP and oxygen. Cells were plated on the same day into a 96-well plate in DMEM/F12 (100 μl) without glucose and supplemented with B27, EGF and bFGF (20 ng/ml), as well as galactose (10 mM) to obtain the ATP amount generated by mitochondrial activity. ATP standards were loaded on the same plate as references. The assay was performed according to the manufacturer’s instructions.

### Oxygen consumption rate and extracellular acidification rate measurements

The functional status of mitochondria in MCAD deficient GSCs 8.11 and NHA was determined by analyzing multiple parameters of oxidative metabolism using the XF96 Extracellular Flux Analyzer (Agilent), which measures extracellular flux changes of oxygen and protons. Cells were plated in XF96-well microplates (30,000 cells per well) in a final volume of 80 μl of DMEM/F12 medium (17.5 mM glucose, 2.5 mM glutamine) supplemented with B27, EGF (20ng/ml), bFGF (20ng/ml). For the mitochondrial stress test, cells were incubated at 37 °C in the absence of CO2 and in 180 μl DMEM-F12 (17.5 mM glucose, 2.5 mM glutamine) supplemented with B27, EGF (20ng/ml), bFGF (20ng/ml of XF-Mito-MEM) per well for 1 hour before the assay. The ports of the sensor cartridge were sequentially loaded with 20 μl per well of the appropriate compound: the ATP coupler oligomycin (Sigma, O4876), the uncoupling agent carbonyl cyanide 4-(trifluoromethoxy) phenylhydrazone (FCCP, Sigma C2920) and the complex I inhibitor rotenone (Sigma, R8875).

### Immunoblotting

Protein lysates were resolved on 4–15% gradient polyacrylamide SDS gels and transferred onto nitrocellulose membranes according to standard procedures. Membranes were incubated with indicated primary antibodies, washed, and probed with HRP-conjugated secondary antibodies. The detection of bands was carried out upon chemiluminescence reaction followed by film exposure. The following primary antibodies were used: Anti-ACADM antibody (ab92461), Anti-Oxoguanine 8 antibody (ab206461), Anti-4 Hydroxynonenal antibody [HNEJ-2] (ab48506), Anti-Vinculin antibody [VIN-54] (ab130007), Anti-GAPDH antibody [EPR6256] (ab128915), Anti-beta Actin antibody (ab8224)

### Transmission electron microscopy (TEM)

TEM was performed at the MDACC High Resolution Electron Microscopy Facility. Samples were fixed with a solution containing 3% glutaraldehyde plus 2% paraformaldehyde in 0.1 M cacodylate buffer, pH 7.3, for 1 hour. After fixation, the samples were washed and treated with 0.1% millipore-filtered cacodylate buffered tannic acid, post-fixed with 1% buffered osmium tetroxide for 30 min, and stained in bloc with 1% millipore-filtered uranyl acetate. The samples were dehydrated in increasing concentrations of ethanol, then infiltrated and embedded in LX-112 medium. The samples were polymerized in a 60 °C oven for 2 days. Ultrathin sections were cut using a Leica Ultracut microtome, stained with uranyl acetate and lead citrate in a Leica EM Stainer, and examined in a JEM 1010 transmission electron microscope (JEOL) at an accelerating voltage of 80 kV. Digital images were obtained using an AMT Imaging System (Advanced Microscopy Techniques Corp).

### Oxidative stress detection

Reduced and oxidized forms of glutathione (GSH and GSSG, respectively) were measured in GSC 8.11 and GSC 6.27 using a GSH/GSSG Ratio Detection Assay Kit (Abcam, ab138881) according to the manufacturer’s protocol. Data from 3 separate experiments were averaged for the results. NADP+ and NADPH levels were measured from GSC 8.11 and 6.27 using a NADP/NADPH Assay Kit (Abcam, ab65349) according to the manufacturer’s instructions. NADP and NADPH levels in total lysate were calculated by comparison with the standard curve.

### Flow Cytometry

Cellular apoptosis was detected using a FITC Annexin V Apoptosis Detection Kit I (BD Biosciences), according to the manufacturer’s instructions. Following virus infection with sh-ACADM or sh-scr and 48 hours of puromycin selection, cells were harvested and re-suspended in cold PBS. Subsequent to centrifugation at 1000 rpm for 5 min at 4°C, the cells were resuspended with 500 μl binding buffer and mixed with 5 μl annexin V-FITC. The cells were subsequently incubated with 5 μl propidium iodide (PI) in the dark at room temperature for 5-15 min. Excitation was at 488 nm and the emission filters used were 515–545 BP (green, FITC) and 620 LP (red, PI). All assays were performed in triplicate.

To detect reactive oxygen species (ROS,) 2×10^5^ cells were stained with Mitotracker Green (100 nM) (Molecular Probes) for 20 min, washed twice and resuspended in PBS. Excitation was at 488 nm and the emission filters used were 515–545 BP (green, FITC). ROS were induced by 4-hydroxynonenal treatment (10 μM) for positive controls. Gating strategies to exclude doublets and dead cells (DAPI) were always employed. After staining, samples were acquired using a BD FACSCantoII flow cytometer. Data were analyzed by BD FACSDiva or FlowJo (Tree Star).

### Immunohistochemistry and immunocytochemistry

For immunohistochemistry (IHC) staining, tumor samples were fixed in 4% formaldehyde for 2 to 4 hours on ice, moved in 70% ethanol for 12 hours, and then embedded in paraffin (Leica ASP300S). After cutting (Leica RM2235), baking and deparaffinization, slides were treated with Citra-Plus Solution (BioGenex) according to specifications. Endogenous peroxidases were inactivated by 3% hydrogen peroxide. Nonspecific signals were blocked using 3% BSA, 10% goat serum and 0.1% Triton. Tumor samples were stained with primary antibodies. ImmPress and ImmPress-AP (Vector Lab) were used as secondary antibodies; Nova RED, Vector BLUE and DAB were used for detection (Vector Lab). Images were captured with a Nikon DS-Fi1 digital camera using a wide-field Nikon EclipseCi microscope. For Oil Red O Lipid staining, T98G were grown on coverslips and GSCs were attached to tissue slides by cytospin. Tumor tissues were 4% PFA fixed, cryoprotected with 30% sucrose, OCT embedded and sectioned (5 μm thick). Oil Red O Lipid staining was performed according to the manufacturer instructions (Abcam, ab150678).

### ACADM shRNAs and sgRNAs

The hairpin RNA interference plasmid for human ACADM, pLKO.1 ACADM TRCN0000028530 (sh-AC1), TRCN0000028509 (sh-AC2) and the scramble control pLKO.1-Puro plasmid (sh-scr) were obtained from Sigma-Aldrich. The sequence of shRNAs are:

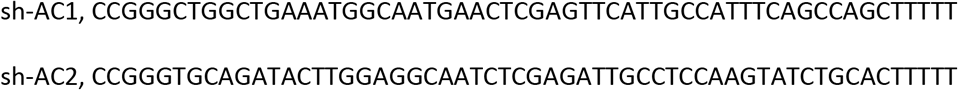

The inducible shRNA was obtained from Cellecta by cloning into a pRSIT16-U6Tet-sh-CMV-TetRep-2A-TagRFP-2A-Puro vector the following hairpin sequence:

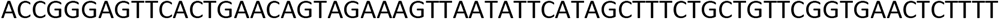

To generate ACADM sgRNAs three pairs of oligonucleotides were designed and used as follows:

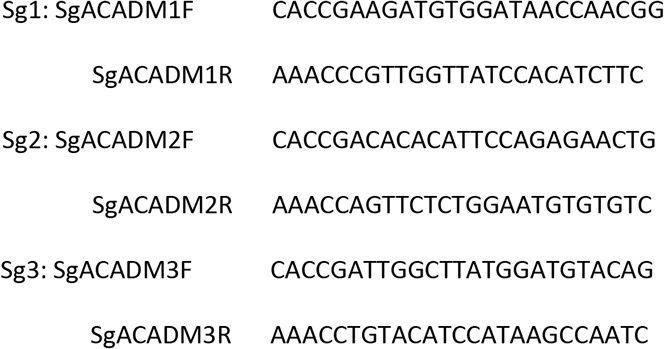

All sgRNAs were validated for their KO efficiency. sgACADM1 was selected for further studies to knockout ACADM. To specifically amplify ACADM in genomic DNA, the following oligonucleotides were used:

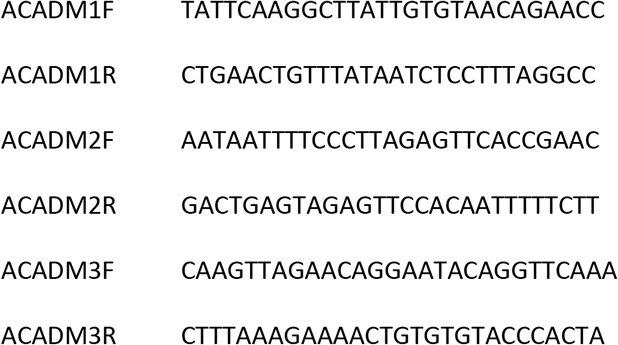

### Lipidomics of total cell extracts and mitochondria

After adding equal volumes of dichloromethane/methanol/PBS (1:1:1), samples were centrifuged (2000 rpm; 5 min) to collect the organic phase. The extraction was repeated twice. After drying using a gentle stream of N2 gas, samples were dissolved in 8 mM ammonium fluoride dichloromethane/isopropyl alcohol/methanol (2:1:1) and sonicated (5 min) before an internal standard was added. Lipids were measured using the shotgun lipidomics by directly infusing a modified Blight-Dyer extract into a SCIEX Triple TOF 5600+ mass spectrometer for MS/MS^ALL analysis as previously described (Padanad et al., 2016). Acquisition was performed once in positive and once in negative ion mode. The phospholipid species were identified based on their characteristic *m/z* value, fragmentation analysis, and precursor ion or neutral loss scans. Using precursor ion scanning techniques, negative precursors of *m/z* 241 and 196 identified parents of phosphatidylinositol, phosphatidylethanolamine, and lysophosphatidylethanolamine. Positive precursors of *m/z* 184 yielded parents of sphingomyelin, phosphatidylcholine, and lysophosphatidylcholine. Neutral loss scans of 141 and 185 yielded parents of phosphatidylethanolamine and phosphatidylserine, respectively. Neutral lipids, including triacylglycerides, diacylglycerides, and cholesterylesters, were identified based on their fatty acid neutral loss in positive ion mode using the Lipid Maps database (http://www.lipidmaps.org/). The same extraction method was used for lipidomics on purified mitochondria. Cardiolipin peaks were identified as their [M-2H]^2-^ ions in the negative-ionization mode by PI scanning of *m/z* 153. The intensity of each peak was normalized to the total lipid signal and to the internal standard. The normalized data relative to each lipid species were summed to give the intensity of each class, which was reported as percentage of all lipids.

### Lentivirus production

Lentivirus was produced by polyethyleneimine (PEI) transfection of 293T cells. Four million cells were transfected with 30 μg of transfer vector, 19 μg of packaging plasmid pCMV-Δ8.74 and 9.5 μg of envelope plasmid pMD2G-VSVG. After 16 hours medium was replaced and viral particles in the medium were collected 72 hours later by ultracentrifugation at 23,000 × g for 3 hours.

### Cells infection

Cells were infected with lentiviral particles expressing shRNAs, inducible shRNAs or sgRNAs as indicated for each experiment. 4 x 10^6^ GSCs were infected with 4 x 10^6^ lentiviral particles and selected for 48 hours with Puromycin (3 μg/ml). In the experiments with NHAs and T98G 1 x 10^6^ cells were infected with 1 x 10^6^ lentiviral particles.

### T7E1 assay

72 hours post transfection or 1 week post lentiviral infection of guide and nuclease, cell genomic DNA was extracted with Qiagen DNeasy blood and tissue kit per manufacture’s protocol. Genomic DNA was used for the template of PCR with NEB Onetaq mastermix per manufacture’s protocol. The product was first denatured at 98°C for 5 minutes, then slowly annealed to 75°C at 1°C /s and eventually to 25°C at 0.1°C /s. 5U T7 Endonuclease I was used for digestion of less than 300 ng annealed product for 30 minutes. The digested product was subjected to 2% TAE gel electrophoresis.

### Fatty Acid Analysis

MCAD silenced or control GSCs were grown for 24 hours after Puromycin selection in complete DMEM-F12. After 24 hours cells were harvested and the pellet were placed at −80°C. Microtubes containing cell pellets were removed from −80°C storage and maintained on wet ice throughout the processing steps. 20 μL of 0.75 μg/ μl D27 myristic acid was spiked in as internal standard (IS). 1mL of methanol was added to each sample, subject to sonication for 2 min, and centrifuged at 4000 rpm for 10 min at 4 °C. Supernatant was transferred into a new tube and dried under N2. For medium samples, 500 μl of sample was spiked with 20 μl D27 myristic acid and prepared as above. Stock solutions of hexanoic acid, octanoic acid and decanoic acid were prepared at 28 μg/μl, 11 μg/μl, 11 μg/μl, respectively, in methanol. The D27 myristic acid IS was spiked into standards at 0.75 μg/μl. Cell lysates, media samples, and standards were then derivatized, as follows. 100 μl of N-methyl-N-trimethylsilyl trifluoroacetamide (MSTFA, Pierce) with 1% trimethylchlorosilane (1% TMCS, Thermo Scientific) was added to react for 30 min in 60°C. 1 μl of sample was injected into GCMS for detection. GC/MS analysis was performed using an Agilent 7890A GC equipped with a 15m DB-5MS+DG capillary column and a Leap CTC PAL ALS as the sample injector. The GC was connected to an Agilent 5975C quadrupole MS operating under positive electron impact ionization at 70eV. All tunings and data acquisition were done with an HP PC with Win 7 professional OS that included the ChemStation E.02.01, PAL Loader 1.1.1, Agilent Pal Control Software Rev A and Pal Object Manager updated firmware. MS tuning parameters are in default settings. GC injection port was set at 250 °C and GC oven temperature was held at 60 °C for 5 min and increased to 220 °C at 10 °C/min, then held for 10 min under constant flow with initial pressure of 10.91 psi. MS source and quadrupole were held at 230 and 159 °C, respectively, and the detector was run in scanning mode, recording ion abundance in the range of 26-600 m/z with solvent delay time of 2 min. The data set was translated into .D format. The extraction was done with Agilent MassHunter WorkStation Software GCMS Quantitative Analysis Version B.07. Identification was performed by searching NIST2011. A one-point calibration was performed in this study and data was normalized by IS to generate the final report.

### ^13^C tracer metabolomics

Cells were grown in complete DMEM-F12 supplemented with^13^C-oleate/ ^13^C-glucose. After 6/12 hours cells were harvested and the pellet Cell pellets were dissolved in 50 μL water/methanol (50:50) and x μL were injected onto a Waters Acquity UPLC BEH TSS C18 column (2.1 x 100mm, 1.7μm) column on an into an Agilent 1260 UHPLC with mobile phase A) consisting of 0.5 mM NH4F and 0.1% formic acid in water; mobile phase (B) consisting of 0.1% formic acid in acetonitrile. Gradient program: mobile phase (B) was held at 1% for 1.5 min, increased to 80% in 15 min, then to 99% in 17 min and held for 2 min before going to initial condition and held for 10 min. The column was at 40 ° C and 3 μl of sample was injected into an Agilent 6520 Accurate-Mass Q-TOF LC/MS. The LC-MS flow rate was 0.2 mL/min. Calibration of TOF MS was achieved through Agilent ESI-Low Concentration Tuning Mix. In negative acquisition mode, key parameters were: mass range 100-1200 da, gas temp 350 °C, fragmentor 150 v, skimmer 65 v, drying Gas 10 l/min, nebulizer flow at 20 psi and Vcap 3500 v, reference ions at 119.0363 and 980.01637 da, ref nebulizer at 20 psi. In positive acquisition mode, key parameters were: key parameters were: mass range 100-1200 da, gas temp 350 °C, fragmentor 150 v, skimmer 65 v, drying Gas 10 l/min, nebulizer flow at 20 psi and Vcap 3500 v, reference ions at 121.050873 and 922.009798 da, ref nebulizer at 20 psi. Agilent Mass Hunter Workstation Software LC/MS Data Acquisition for 6200 series TOF/6500 series Q-QTOF Version B.06.01 was used for calibration and data acquisition.

### Acyl-carnitine profiling

GSC 8.11 were grown for 24 hours in complete DMEM-F12 supplemented with ^13^C-oleate. After 24 hours cells were harvested and the pellet were placed at −80°C. Microtubes containing cell pellets were removed from −80°C storage and maintained on wet ice throughout the processing steps. To initiate protein precipitation, 0.3 mL of a chilled mixture of chilled mixture of methanol and chloroform (8:2) (EMD, Billerica, MA) was added to each sample, the mixture was vortexed briefly, and allowed to incubate on ice for 10 mins. Post-incubation, the vortex step was repeated, and samples centrifuged at 14,000 RPM, for 10 mins in 4°C. Post-centrifugation, 100 μL of supernatant was transferred to an autosampler vial for LC-MS analysis. From the remaining supernatant from each sample, a small aliquot was transferred to a new microtube to create a pooled sample for quality control purposes.

LC-MS analysis was performed on an Agilent system consisting of a 1290 UPLC module coupled with a 6490 QqQ mass spectrometer (Agilent Technologies, CA, USA.) A 1 μL injection of acyl carnitine metabolites were separated on an Acquity HSS-T3 1.8 μM, 2.1 x 50 mm column (Waters, Milford, MA) maintained at 40°C, using 10 mM ammonium acetate in water, adjusted to pH 9.9 with ammonium hydroxide, as mobile phase A, and acetonitrile as mobile phase B. The flow rate was 0.25 mL/min and the gradient was linear 0% to 80% A over 7 mins, then 80 to 100% over 1.5 mins, followed by isocratic elution at 100% A for 5 minutes. The system was returned to starting conditions for 3 mins to allow for column re-equilibration before injecting another sample. The mass spectrometer was operated in ESI-mode with the following instrument settings: Gas temp: 275°C, flow: 15 l/min, nebulizer: 35 psi, capillary 3500 V, sheath gas 250°C, and sheath gas flow 11 l/min. The ion funnel high/low pressure RF settings were: 150/60 V respectively. Acyl carnitine transitions were monitored for the 85 Da product ion that is common to each carnitine species. Data Analysis: Metabolites were identified by matching the retention time and mass (+/- 10 ppm) to authentic standards. Isotope peak areas were integrated using MassHunter Quantitative Analysis vB.07.00 (Agilent Technologies, Santa Clara, CA.) Peak areas were corrected for natural isotope abundance using an in-house written software package based on the method of (Fernandez et al., 1996), and the residual isotope signal was reported. Data were normalized to cell protein content prior to analysis of metabolite fluxes for Central Carbon, Acyl carnitine and Fatty acid metabolites.

## SUPPLEMENTAL FIGURE LEGENDS

**Figure S1**: **Details of metabolome screen in human GSCs.**

**a.** KEGG pathway composition of the metabolome shRNA library. Dark gray bars represent the genes covered by the screen within the total genes that constitute the pathway (light gray). **b.** List of genomic events and driver mutations that characterized the GBM models used for the screen (GSC 8.11 and 6.27). **c.** Kaplan-Meier survival analysis of animals with intracranial tumors generated from injecting 1 x 10^6^ GSC 8.11 or 6.27 cells. **d.,e.** Size-normalized counts (distribution) of *in vivo* metabolome shRNA screens in GSC models for GSC 8.11 and GSC 6.27. Replicates of reference GSC populations upon infection are highlighted in red and blue. Three independent *in vivo* tumors are highlighted in green. **f.** Scatter plot comparing relative depletion of *in vivo* metabolome shRNAs (RSA logP) in two independent GSC models (y axis = GSC 8.11, x axis = GSC 6.27). Unique gene hits in each GSC model are highlighted in red (GSC 8.11) and blue (GSC 6.27). Common gene hits across the two GSC models are highlighted in green. **g.** Relative abundance of *ACADM* in brain cells expressed as Fragments Per Kilobase Of Exon Per Million Fragments Mapped (FPKM, data from RNAseq database from Ben Barres lab at Stanford (available at https://web.stanford.edu)

**Figure S2: Fatty acid oxidation fuels oxidative metabolism in GBM cells.**

**a.** Representative immunoblot of MCAD expression in NHAs in comparison with different GSCs models. GAPDH served as loading control. **b.** Representative immunoblot of FAO enzymes expression in NHAs in comparison with different GSCs models. Vinculin and GAPDH served as loading control. **c.** Isotopologue patterns for incorporation of ^13^C-labeled oleate into TCA cycle intermediates, as measured by LC-MS in NHAs and GSCs in basal conditions in GSCs 8.11 and 272. Cells were cultured with ^13^C oleate for 6 hours prior to sample collection. n = 4 biological replicates, error = +/- SD. \**P*= 0.0129, \*\**P*= 0.0014, \*\*\*\**P* <0.0001.

**Figure S3: MCAD ablation dramatically affects GSC proliferation *in vitro*.**

**a.** Representative immunoblot (left) and quantification (right) of MCAD expression in GSC 8.11 infected with *ACADM*-targeting or control shRNA. Vinculin served as a loading control. **b.** Growth curve of GSCs 6.27, 272, and 7.11 upon shRNA-mediated MCAD knockdown. Values represent the mean ± SD of three independent experiments; *P* values were generated using one-way ANOVA test. GSCs 6.27 \**P*< 0.04; \*\**P* = 0.0051. GSCs 272, * *P* = 0.0132, ** *P* = 0.0078. GSCs 7.11, * *P* = 0.034; ** *P* ≤ 0.004. **c.** (Top) Analysis of Cas9 activity by T7 endonuclease I assay. GSCs were transfected and total genomic DNA was isolated 72 h post transfection for subsequent T7EI cleavage. (Bottom) MCAD knockdown/knockout efficiency of shRNAs/sgRNAs determined by immunoblot. Vinculin served as loading control. **d.** Growth curve of GSC 8.11 cells carrying three independent sgRNA-mediated *ACADM* deletions, compared to a sgRNA control. Day 0 was defined as starting 48 hours post-puromycin selection. Values represent the mean ± SD of three independent experiments. *P* values were generated using one-way ANOVA test. * *P* =0.0146, ** *P* ≤ 0.001. **e.** Growth curve of GSCs 6.27, 272, and 7.11 upon sgRNA-mediated *ACADM* knockout. Values represent the mean ± SD of three independent experiments; *P* values were generated using one-way ANOVA test. GSCs 6.27 * *P* < 0.03; GSCs 272, * *P* ≤ 0.02. GSCs 7.11, * *P* ≤ 0.04. Bottom: Representative immunoblot of MCAD expression in GSC 6.27, 272, 7.11 infected with *ACADM*-targeting or control sgRNA. Vinculin or GAPDH served as a loading control.

**Figure S4: MCAD inhibition triggers apoptotic cell death and attenuates proliferative capacity and sphere-forming efficiency in GSCs *in vitro*.**

**a.** Representative scatter plot showing Annexin V and DAPI staining in GSC 8.11 cells upon MCAD knockdown (7 days post-puromycin selection) as measured by flow cytometry (Left). Quantification of apoptosis in GSC 6.27 infected with anti-*ACADM* shRNA/shRNA by Annexin V (Right). Staining was evaluated by flow cytometry at 72 hours and 7 days after puromycin selection. Values represent the mean ± SD of three or four independent experiments; *P* values were generated using one-way ANOVA test. \**P* < 0.05. **b.** Quantification of apoptosis in GSC 272, 7.11 infected with anti-*ACADM* shRNA/sgRNA by Annexin V. Staining was evaluated by flow cytometry at day 7 after puromycin selection. Values represent the mean ± SD of four independent experiments; *P* values were generated using one-way ANOVA test. 272 \**P* < 0.03, 7.11 \**P* ≤ 0.04. **c.** Cell viability assessed by Trypan Blue exclusion of GSC 6.27 treated with indicated concentration of SPA for 96 hours. Mean values ± SD of three biologically independent replicates. *P* values were generated using one-way ANOVA. \*\**P* = 0.006 **d.** Dot plot showing GSCs 6.27 sphere formation efficiency (SFE) upon SPA treatment at 48 hours at different concentrations. DMSO was used as control. Values represent the mean ± SD of four independent experiments; *P* values were generated using one-way ANOVA test. \**P* = 0.0327.

**Figure S5: MCAD ablation/inhibition impacts tumorigenicity of GSCs in vivo.**

**a.** Luciferase imaging of representative tumors 30 days (GSC 8.11) or 60 days (GSC 6.27) post-implantation. **b.** Quantification of tumor progression after implantation of GSCs 6.27 as measured by MR volumetry (n = 8 mice per group). Data represent mean ± SEM. *P* values were generated using one-way ANOVA. \**P* =0.0121, \*\**P* = 0.0047. **c.** Kaplan-Meier survival analysis upon *in vivo* implantation of GSCs 6.27, 272 and 7.11. For sh-ctrl, sh-*ACADM1*, sh-*ACADM2*, n = 8 mice per group. *P* values were generated using log-rank test. GSC 272 \*\*\**P* = 0.0007; GSC 6.27 and 7.11 **** *P* < 0.0001. **d.** Effect of shRNA-mediated knockdown on tumor growth in GSC 8.11 cells with induced (doxycycline) MCAD depletion 20 days after subcutaneous implantation. Day 0 indicates the beginning of doxycycline treatment (n = 5 mice per group). Values represent the mean ± SD of three independent experiments; *P* values were generated using one-way ANOVA test. \**P* = 0.036, \*\**P* = 0.003; **e.** Representative images (left) of NHAs upon MCAD depletion at 7 days from the beginning of the growth curve shown in Fig. 2h and corresponding immunoblot (right). Vinculin served as loading control.

**Figure S6. MCAD depletion alters the bioenergetics profile of GSCs.**

**a.** Transmission electron microscopy images of mitochondria in GSC 6.27 upon MCAD silencing (magnification x15,000 and 50,000; scale bar= 1 μm, 300 nm). **b.** Analysis of basal respiration rate, maximal respiratory capacity, and ATP-linked respiration generated via metabolic flux assay. Rates are quantified by normalization of OCR level to the total protein OD values. *P* values were generated using one-way ANOVA test \*\**P* = 0.001, **** *P* = 0.0001. Results represent cumulative analyses of multiple independent biological replicates from three independent experiments (n≥4 for each condition per experiment). Data extracted from the OCR traces in Fig. 3C, D. **c.** Isotopologue patterns for incorporation of ^13^C-labeled glucose into TCA cycle intermediates, as measured by LC-MS in GSC 8.11 infected with anti-*ACADM* or control shRNA. Cells were cultured with ^13^C glucose for 24 hours prior to sample collection. N = 3 biological replicates, error bars = +/- SD. **d., e.** 96-h growth curve of GSC 8.11 cells upon *ACADM* silencing with shRNA/sgRNA cultured in DMEM-F12 media supplemented with acetate (2 mM) (d) or linoleic acid (0.1 mg/ml) (e). Day 0 was defined as 48 hours of puromycin selection. Values represent the mean ± SD of three independent experiments. *P* values were generated using one-way ANOVA. In **d**: \**P* = 0.011 ** *P* = 0.003. In **e**: \*\**P* ≤ 0.0033.

**Figure S7. Toxic lipid species are causally correlated with ROS increase and cell death in MCAD-silenced GSCs.**

**a.** Oil Red O staining of GSC 8.11 cells after 72-hour treatment with SPA. DMSO was used as control, scale bar 50 μm. **b.** Cell viability as assessed by Trypan Blue exclusion in *ACADM* wild type or null T98G cells grown in DMEM supplemented with FBS or in DMEM supplemented with LPDS 7 days after puromycin selection. Data represent the mean ± SEM of four biologically independent replicates. *P* values were generated using one-way ANOVA. \*\**P* = 0.0011. **c.** LC-MS acyl-carnitine labeling profile of growth medium and whole-cell extracts from GSC 8.11 infected with anti-*ACADM* or non-targeting shRNAs cultured with ^13^C oleate for 24 hours prior to sample collection. N = 3 biological replicates, error bars represent ± SD. *P* values were generated using one-way ANOVA. \**P* = 0.04, ** *P* = 0.0032 *** *P* = 0.0002, **** *P* <0.0001. **d.** Cell viability assessed by Trypan Blue exclusion of GSC 8.11 cells treated with 400 μM butyrate (C4), hexanoate (C6), octanoate (C8), decanoate (C10), dodecanoate (C12), palmitic acid (C16), or vehicle control for 48 h. Data represent mean ± SEM of four biologically independent replicates. *P* values were generated using one-way ANOVA. \*\**P* < 0.006; \*\*\**P* < 0.0003; \*\*\*\**P* <0.0001. **e.** Quantification by flow cytometry of ROS production in GSC 8.11 cells treated with C10 for 48 h measured by CellROX Green (left) and representative fluorescence (right). Data represent mean ± SEM of three biologically independent replicates. *P* values were generated using two-tailed t-test. \*\**P* = 0.0082. **f.** Viability of GSC 8.11 cells grown for 48 h in medium supplemented with 400 or 800 μM C10 and in the presence or absence of GSH-ethyl ester (GSH-EE) assessed by Trypan Blue exclusion. Data represent mean ± SEM of four biologically independent replicates. *P* values were generated using one-way ANOVA. \**P*= 0.0163; *** *P*= 0.0005. **g.** Quantification of cell viability as assessed by Annexin V in *ACADM* wild type or null GSCs 272 cells grown in DMEM supplemented with ferrostatin-1 (0.5 μM) 7 days after puromycin selection. Data represent the mean ± SEM of four biologically independent replicates. *P* values were generated using one-way ANOVA. **** *P* <0.0001.

**Figure S8. MCAD knockdown triggers ROS-related damage in GSCs *in vitro* and *in vivo*.**

**a,b,c.** Quantification of ROS production (mean of fluorescence intensity) as measured by CellROX Green in GSC 6.27 (a), 272 (b) and 7.11 (c) cells harboring anti-*ACADM* or non-targeting shRNA. Values represent the mean ± SD of three or four independent experiments. *P* values were generated using one-way ANOVA test. (a) \**P* = 0.025, *** *P* = 0.0009, (b) ** *P* = 0.007, **** *P* < 0.0001, (c) \**P* < 0.03. d. Immunostaining for MCAD of xenograft tumor samples derived from GSC 8.11 harboring doxycycline-inducible anti-*ACADM* or non-targeting shRNAs (shown in Fig. 5E and ED Fig. 5D) and relative quantification. Quantification was performed with ImageJ software, four/five images per condition, where each condition was represented by three biological replicates. Values are expressed as mean ± SD; *P* values were generated using one-way ANOVA test. *** *P* = 0.0002. e. Growth curves of GSC 6.27, 272 and 7.11 cells infected with sh-*ACADM* or non-targeting shRNAs grown in the presence or absence of GSH-EE. Values are expressed as mean ± SD; *P* values were generated using one-way ANOVA test. GSC 6.27, ** *P<* 0.007; GSC 272 \**P* = 0.03, ** *P* ≤ 0.004; GSC 7.11 \**P* ≤ 0.04. f. Quantification of the number of spheres formed by GSC 8.11 and GSC 6.27 harboring sh-*ACADM* or non-targeting shRNAs grown for 7 days (for 8.11) or 72 hours and 7 days (for 6.27) in the presence or absence of GSH-EE as indicated. Values represent the number of spheres per field expressed as mean ± SD. *P* values were generated using one-way ANOVA test. \**P* ≤ 0.04. g. Quantification of ROS via CellRox Green staining intensity by flow cytometry in *ACADM* wild type or null GSC 8.11 cells grown in FBS or LPDS. Mean values ± SD of three biologically independent replicates. *P* values were generated using one-way ANOVA. **** *P* = 0.0001. MFI= Mean of Fluorescence Intensity.

