## Supplemental Figures for "Medium-chain acyl-CoA dehydrogenase, a gatekeeper of mitochondrial function in glioblastoma multiforme"

a

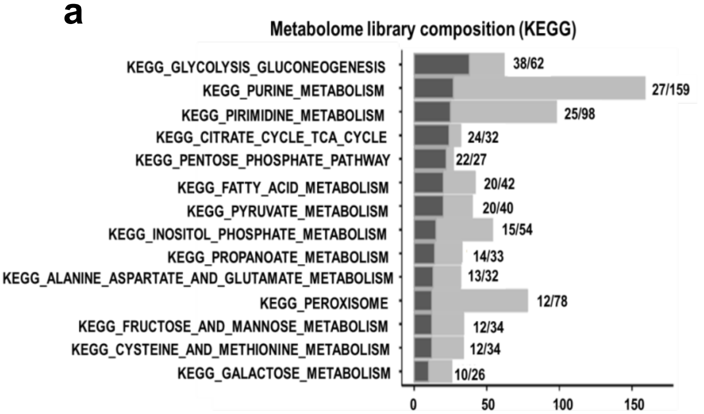


b

| | Subtype: | GSC 8.11 Proneural | GSC 6.27 Classical |
|---|---|---|---|
| Copy Number | EGFR | Neutral | Neutral |
| | PTEN | Neutral | Loss |
| | CDK4 | Neutral | Neutral |
| | RB1 | Loss | Neutral |
| | CDKN2A/B | Loss | Loss |
| | MDM2 | Neutral | Neutral |
| | MDM4 | Neutral | Neutral |
| | PDGFRA | Neutral | Neutral |
| Mutation | IDH1/2 | WT | WT |
| | EGFR | WT | WT |
| | TP53 | Missense | WT |
| | RB1 | Missense | WT |
| | PTEN | Splice site | WT |
| | NF1 | WT | WT |
| | PDGFRA | WT | WT |

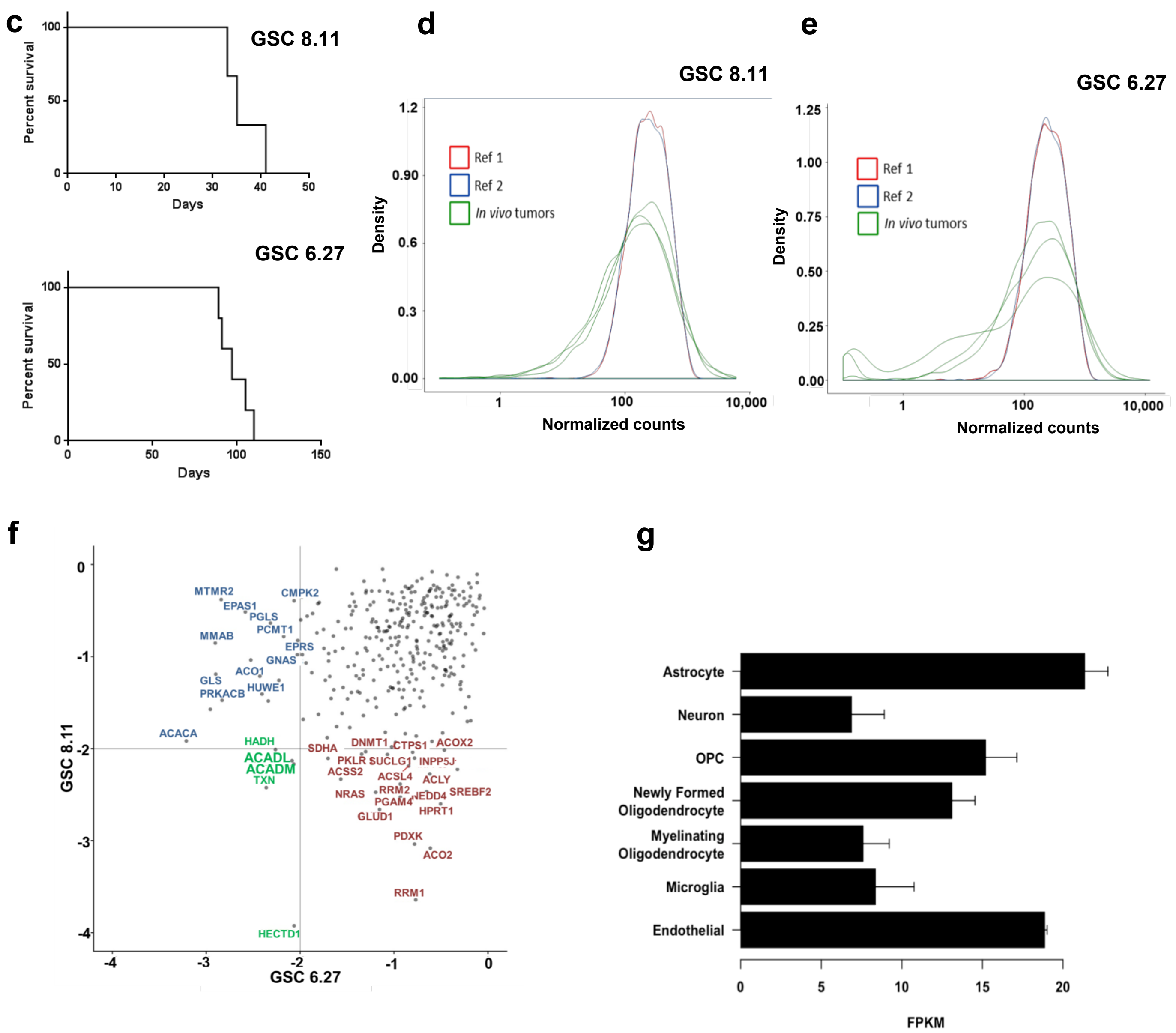


Figure S1

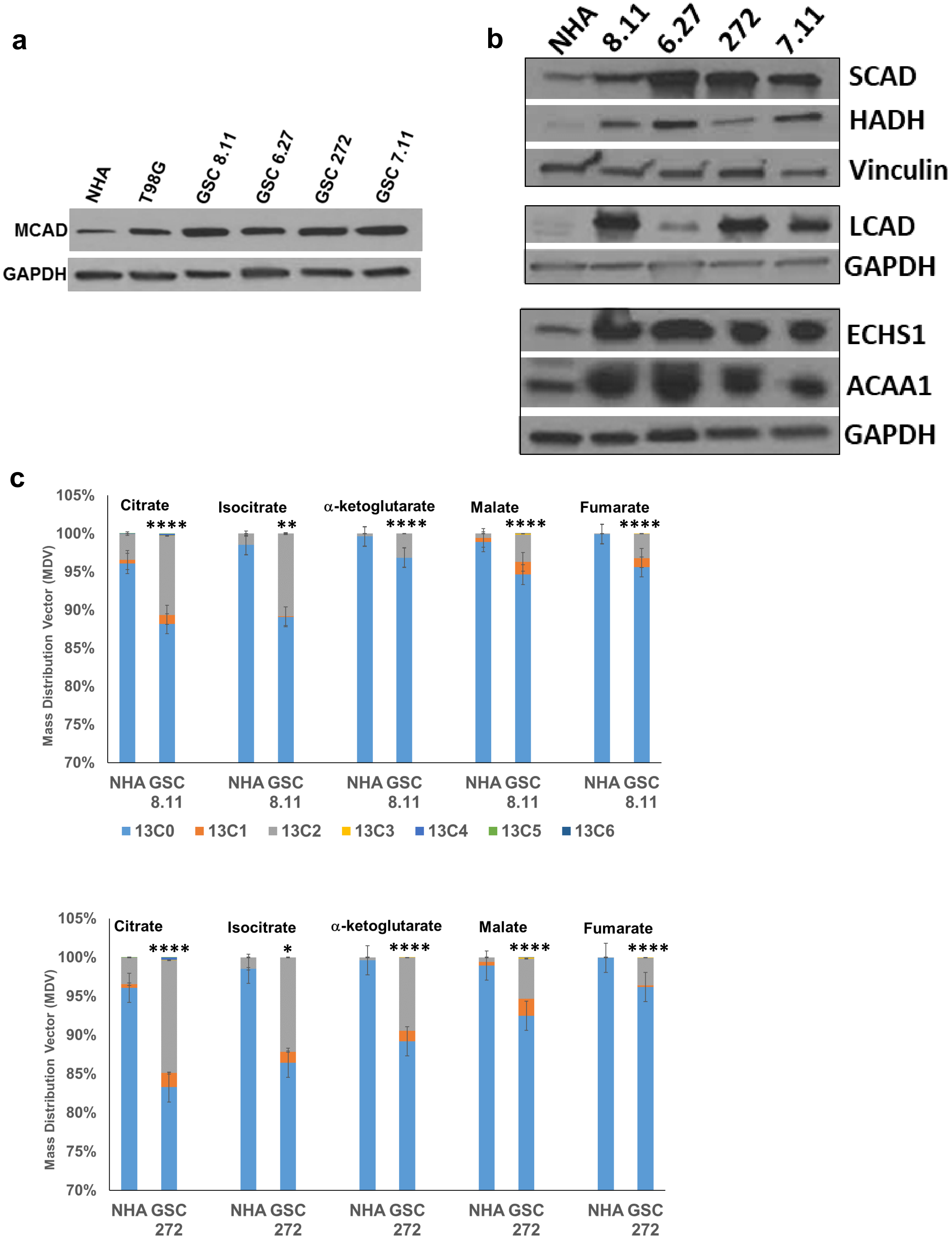
a
NHA
T98G
GSC 8.11
GSC 6.27
GSC 272
GSC 7.11
MCAD
GAPDH
b
NHA
8.11
6.27
272
7.11
SCAD
HADH
Vinculin
LCAD
GAPDH
ECHS1
ACAA1
GAPDH
c
Mass Distribution Vector (MDV)
105%
100%
95%
90%
85%
80%
75%
70%
Citrate
Isocitrate
α-ketoglutarate
Malate
Fumarate
****
**
****
****
****
NHA GSC 8.11
13C0
13C1
13C2
13C3
13C4
13C5
13C6
Mass Distribution Vector (MDV)
105%
100%
95%
90%
85%
80%
75%
70%
Citrate
Isocitrate
α-ketoglutarate
Malate
Fumarate
****
*
****
****
****
NHA GSC 272
13C0
13C1
13C2
13C3
13C4
13C5
13C6

Figure S2

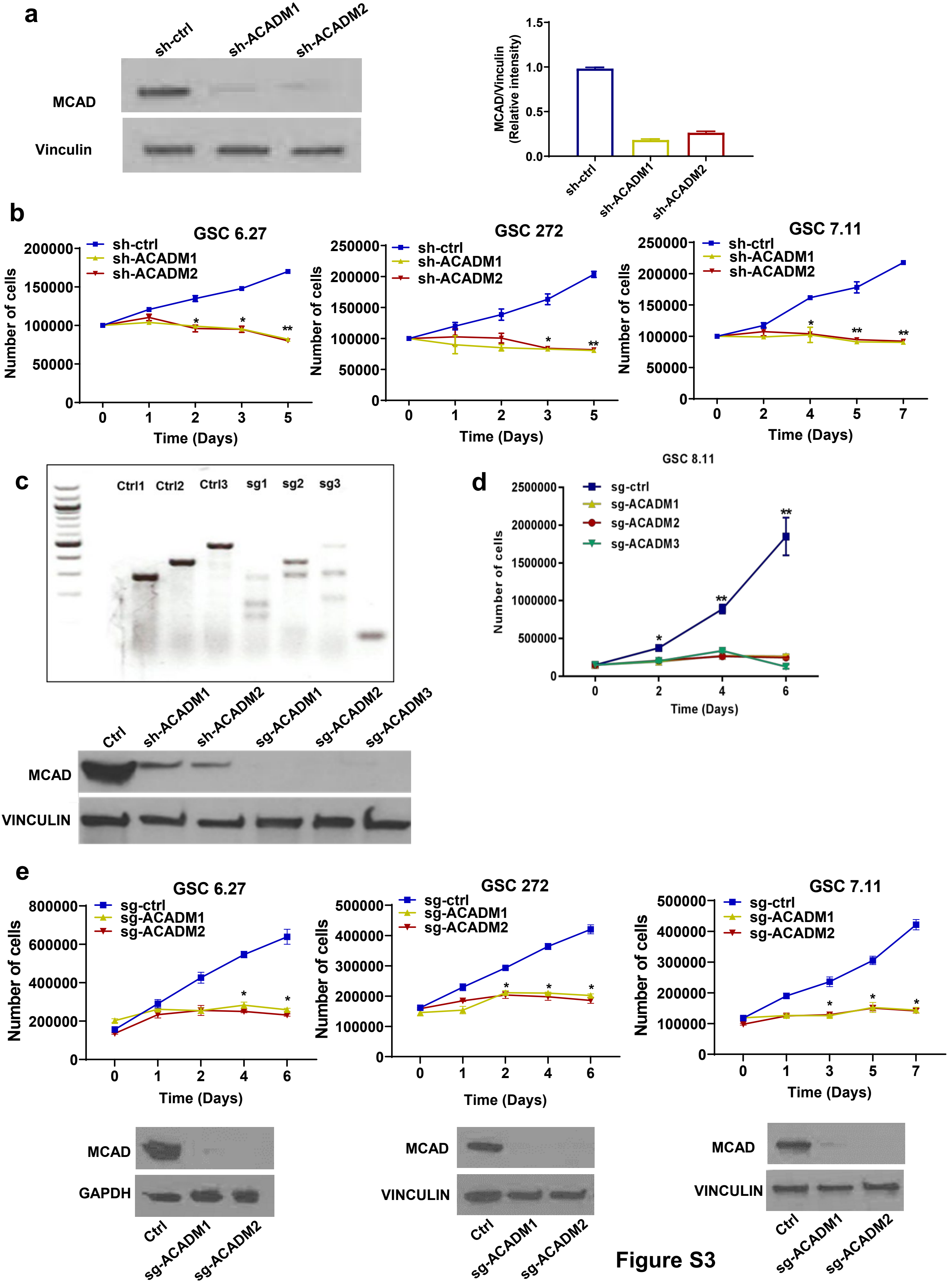
a
sh-ctrl
sh-ACADM1
sh-ACADM2
MCAD
Vinculin
MCAD/Vinculin (Relative intensity)
1.5
1.0
0.5
0.0
b
GSC 6.27
GSC 272
GSC 7.11
sh-ctrl
sh-ACADM1
sh-ACADM2
Number of cells
Time (Days)
c
Ctrl1
Ctrl2
Ctrl3
sg1
sg2
sg3
Ctrl
sh-ACADM1
sh-ACADM2
sg-ACADM1
sg-ACADM2
sg-ACADM3
MCAD
VINCULIN
d
GSC 8.11
sg-ctrl
sg-ACADM1
sg-ACADM2
sg-ACADM3
Number of cells
Time (Days)
e
GSC 6.27
GSC 272
GSC 7.11
sg-ctrl
sg-ACADM1
sg-ACADM2
Number of cells
Time (Days)
MCAD
GAPDH
VINCULIN
Ctrl
sg-ACADM1
sg-ACADM2

Figure S3

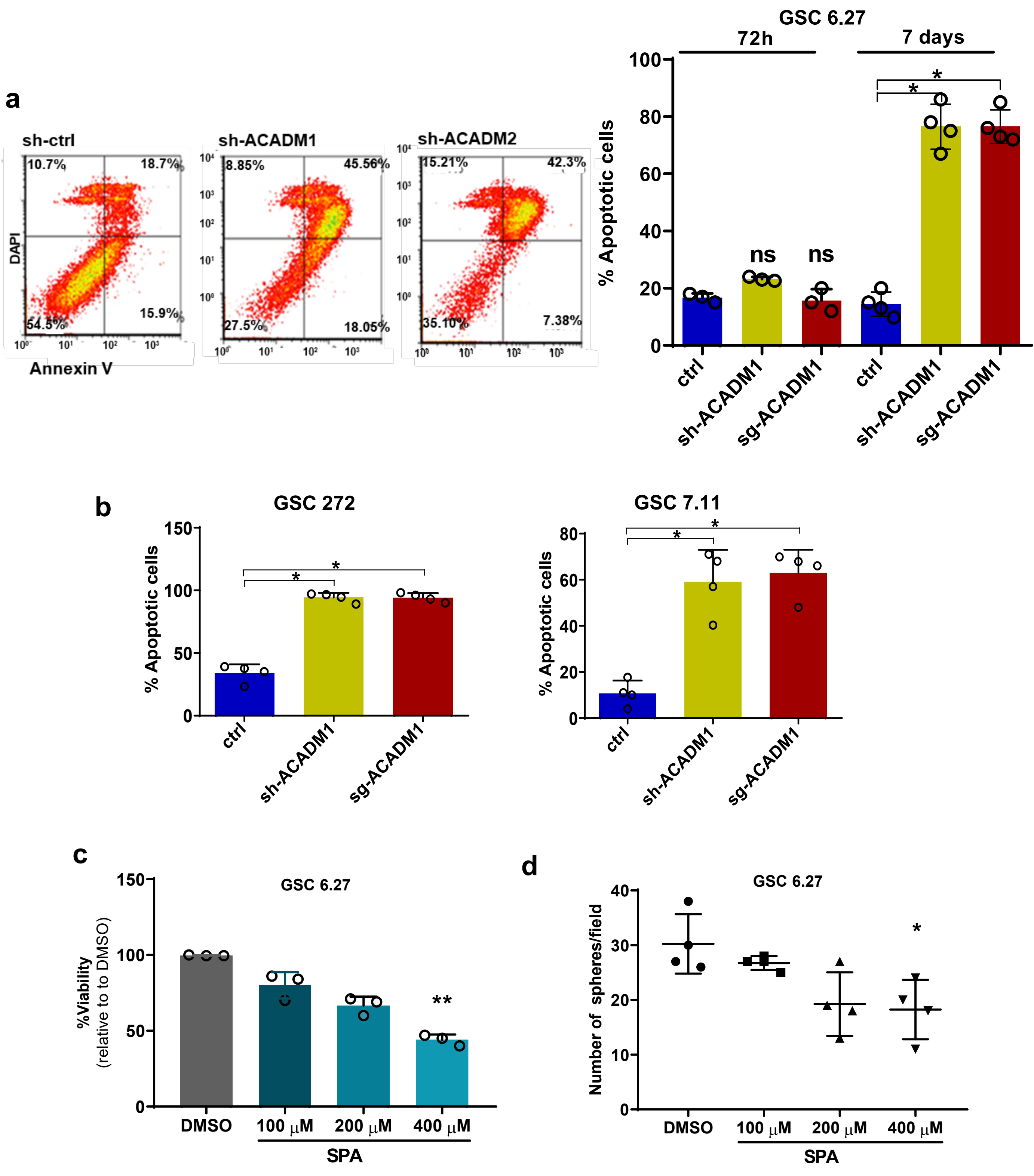

a
sh-ctrl
sh-ACADM1
sh-ACADM2
DAPI
Annexin V
10.7%
18.7%
54.5%
15.9%
8.85%
45.56%
27.5%
18.05%
15.21%
42.3%
35.10%
7.38%
GSC 6.27
72h
7 days
% Apoptotic cells
ns
ns
ctrl
sh-ACADM1
sg-ACADM1
b
GSC 272
GSC 7.11
c
GSC 6.27
%Viability
(relative to to DMSO)
DMSO
100 μM
200 μM
400 μM
SPA
d
GSC 6.27
Number of spheres/field


Figure S4

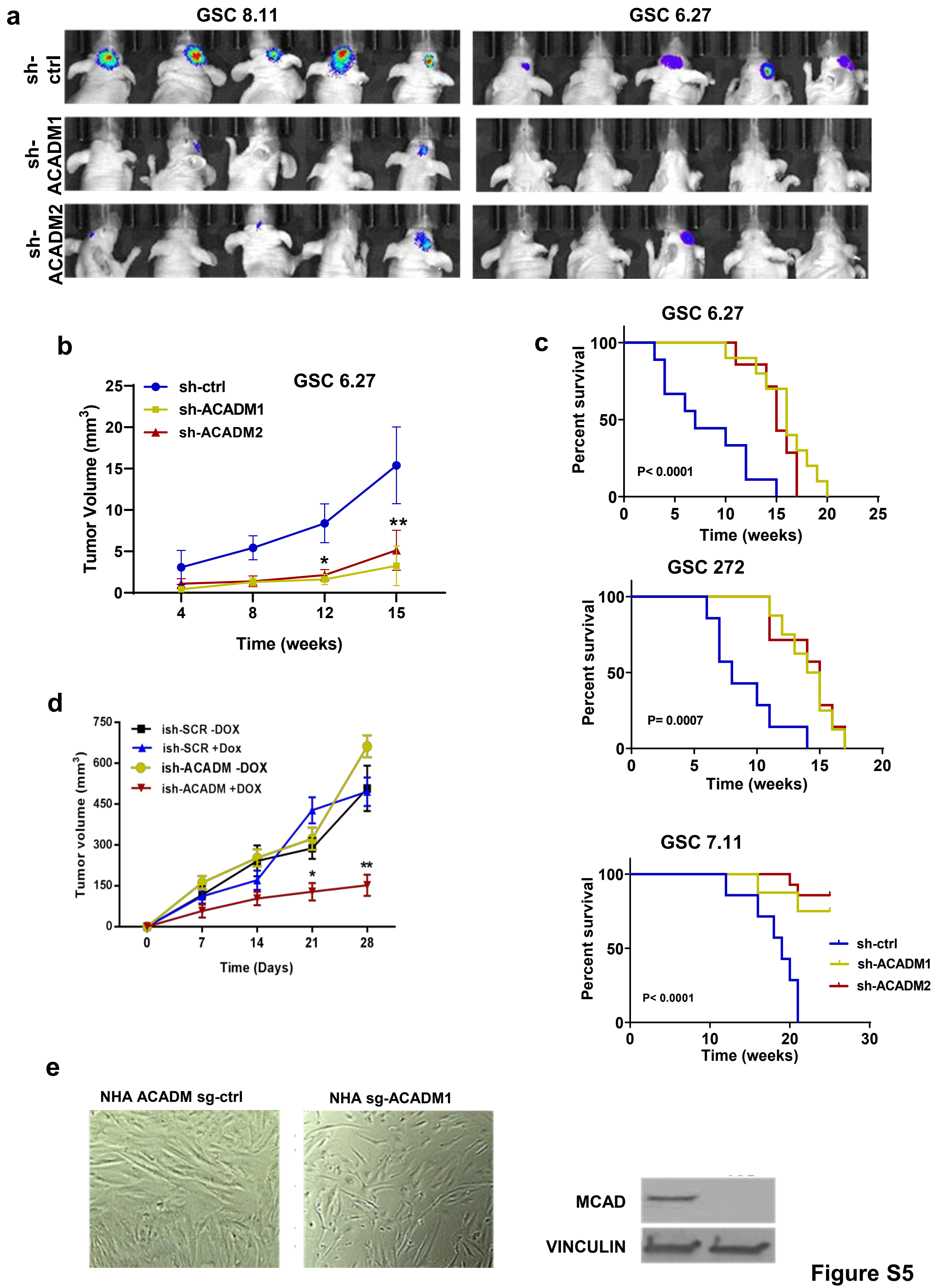

a
GSC 8.11
GSC 6.27
sh-ctrl
sh-ACADM1
sh-ACADM2
b
GSC 6.27
sh-ctrl
sh-ACADM1
sh-ACADM2
Tumor Volume (mm3)
Time (weeks)
c
GSC 6.27
Percent survival
P< 0.0001
Time (weeks)
GSC 272
P= 0.0007
GSC 7.11
P< 0.0001
sh-ctrl
sh-ACADM1
sh-ACADM2
d
ish-SCR -DOX
ish-SCR +Dox
ish-ACADM -DOX
ish-ACADM +DOX
Tumor volume (mm3)
Time (Days)
e
NHA ACADM sg-ctrl
NHA sg-ACADM1
MCAD
VINCULIN


Figure S5

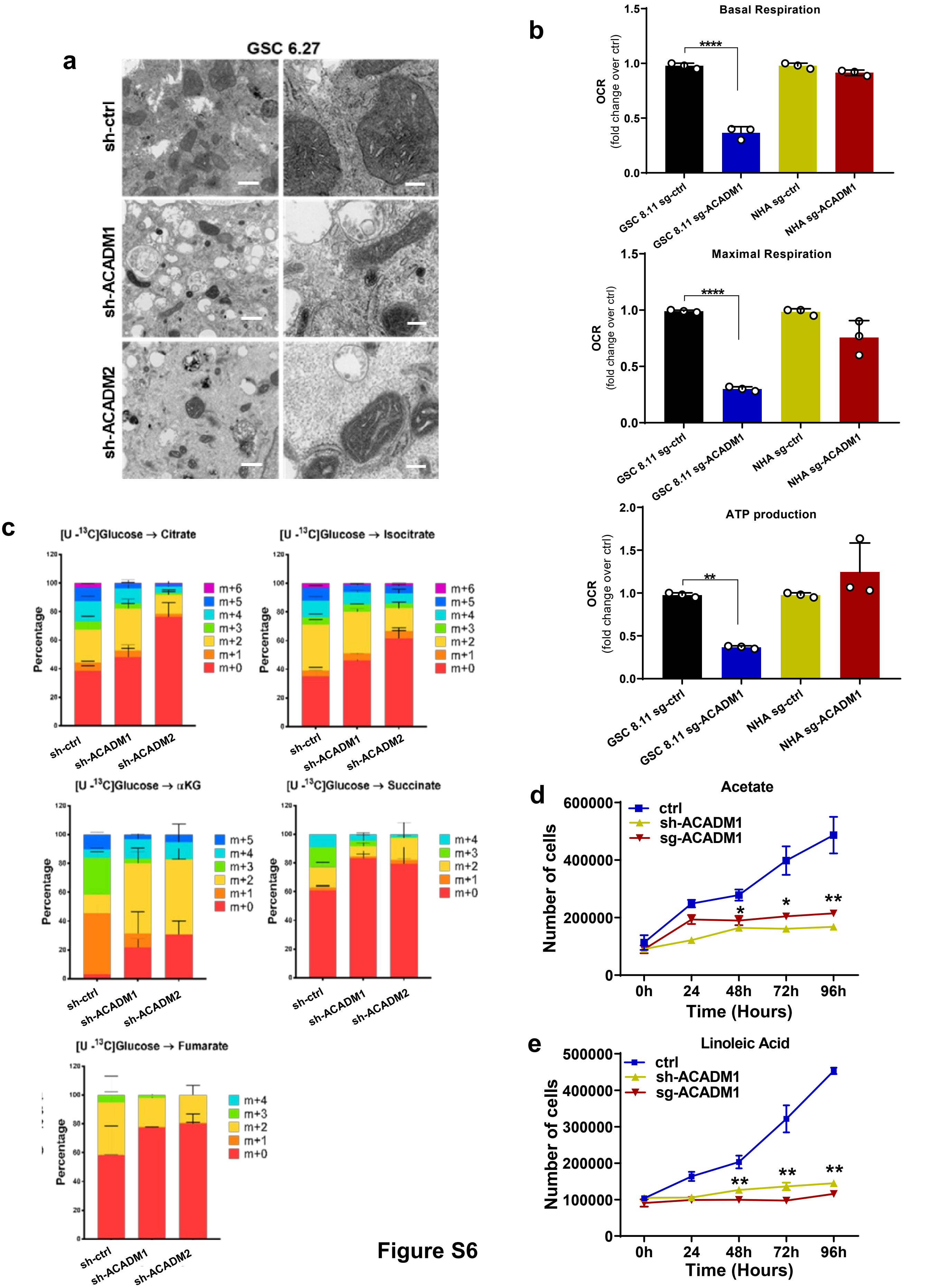
a
GSC 6.27
sh-ctrl
sh-ACADM1
sh-ACADM2
b
Basal Respiration
****
OCR
(fold change over ctrl)
1.5
1.0
0.5
0.0
GSC 8.11 sg-ctrl
GSC 8.11 sg-ACADM1
NHA sg-ctrl
NHA sg-ACADM1
Maximal Respiration
****
ATP production
**
2.0
c
[U -13C]Glucose → Citrate
[U -13C]Glucose → Isocitrate
[U -13C]Glucose → αKG
[U -13C]Glucose → Succinate
[U -13C]Glucose → Fumarate
Percentage
m+6
m+5
m+4
m+3
m+2
m+1
m+0
sh-ctrl
sh-ACADM1
sh-ACADM2
d
Acetate
ctrl
sh-ACADM1
sg-ACADM1
Number of cells
600000
400000
200000
0
0h
24
48h
72h
96h
Time (Hours)
*
*
**
e
Linoleic Acid
500000
400000
300000
200000
100000
**
**
**

Figure S6

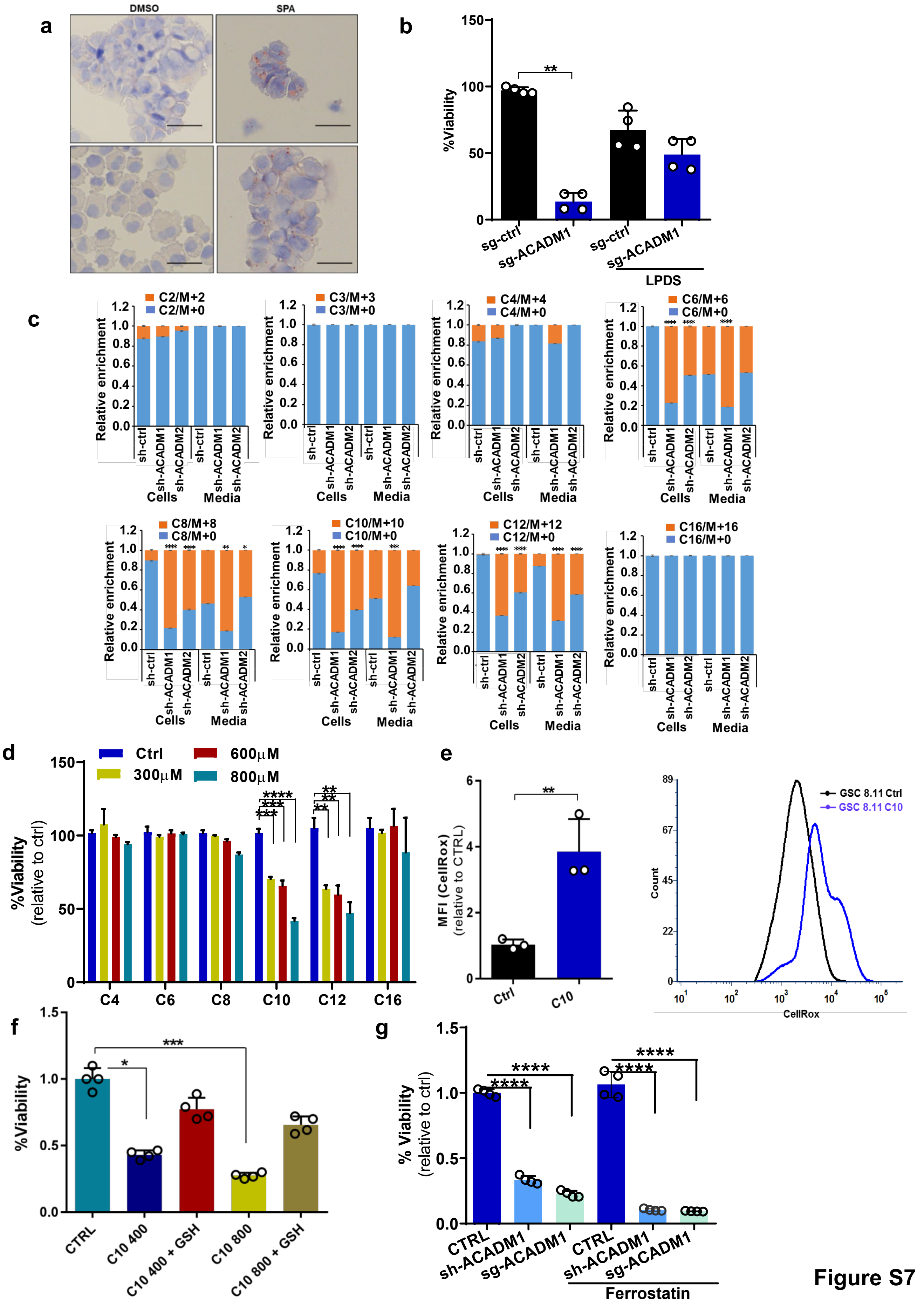
a
DMSO
SPA
b
%Viability
**
sg-ctrl
sg-ACADM1
sg-ctrl
sg-ACADM1
LPDS
c
C2/M+2
C2/M+0
C3/M+3
C3/M+0
C4/M+4
C4/M+0
C6/M+6
C6/M+0
C8/M+8
C8/M+0
C10/M+10
C10/M+0
C12/M+12
C12/M+0
C16/M+16
C16/M+0
Relative enrichment
sh-ctrl
sh-ACADM1
sh-ACADM2
Cells
Media
d
Ctrl
600μM
300μM
800μM
%Viability
(relative to ctrl)
C4
C6
C8
C10
C12
C16
e
MFI (CellRox)
(relative to CTRL)
Ctrl
C10
GSC 8.11 Ctrl
GSC 8.11 C10
Count
CellRox
f
%Viability
CTRL
C10 400
C10 400 + GSH
C10 800
C10 800 + GSH
g
% Viability
(relative to ctrl)
CTRL
sh-ACADM1
sg-ACADM1
Ferrostatin

Figure S7

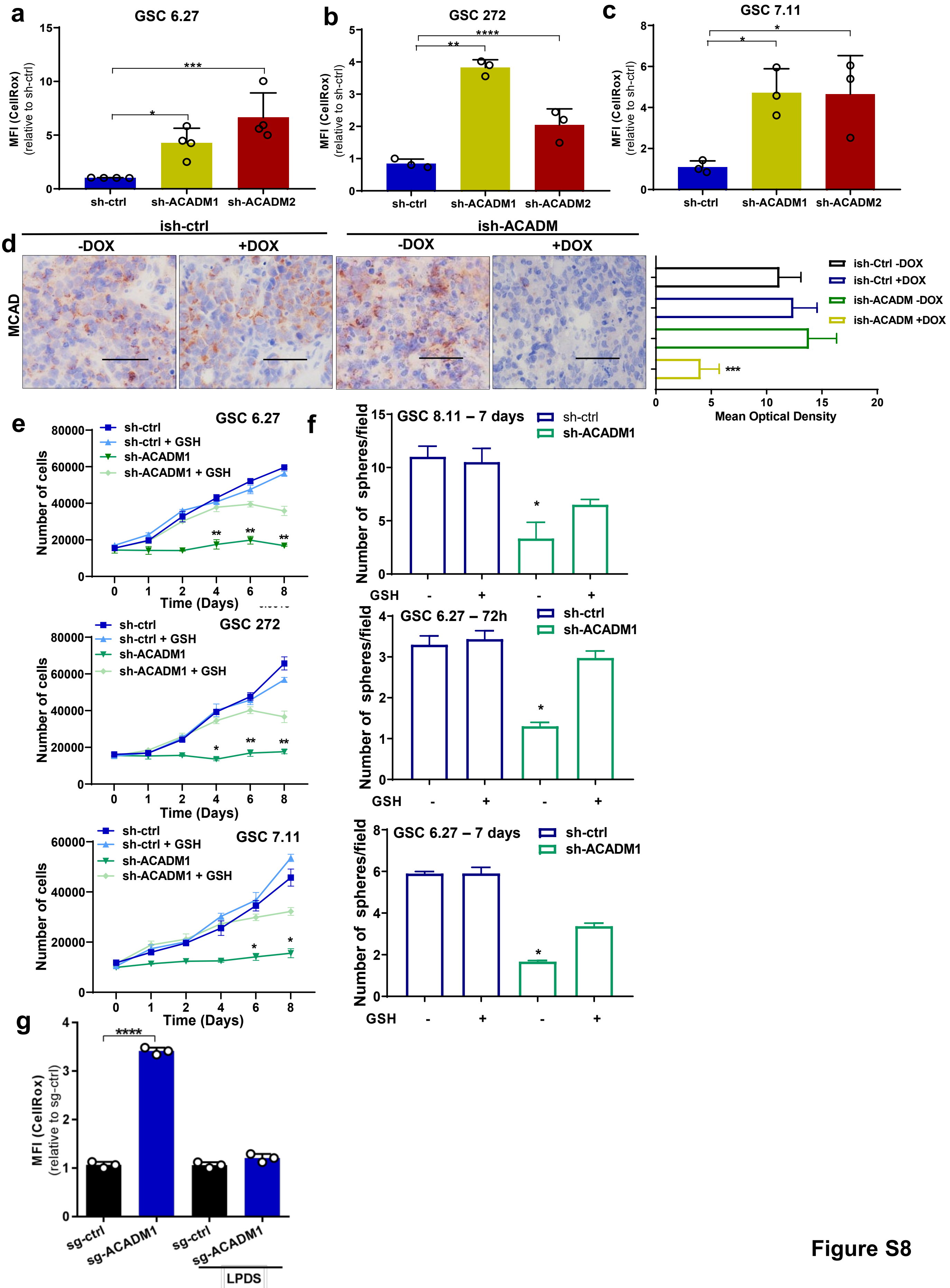
a
GSC 6.27
MFI (CellRox)
(relative to sh-ctrl)
sh-ctrl
sh-ACADM1
sh-ACADM2
*
***
b
GSC 272
MFI (CellRox)
(relative to sh-ctrl)
sh-ctrl
sh-ACADM1
sh-ACADM2
**
****
c
GSC 7.11
MFI (CellRox)
(relative to sh-ctrl)
sh-ctrl
sh-ACADM1
sh-ACADM2
*
*
d
ish-ctrl
ish-ACADM
-DOX
+DOX
-DOX
+DOX
MCAD
ish-Ctrl -DOX
ish-Ctrl +DOX
ish-ACADM -DOX
ish-ACADM +DOX
***
Mean Optical Density
e
GSC 6.27
sh-ctrl
sh-ctrl + GSH
sh-ACADM1
sh-ACADM1 + GSH
Number of cells
Time (Days)
GSC 272
sh-ctrl
sh-ctrl + GSH
sh-ACADM1
sh-ACADM1 + GSH
Number of cells
Time (Days)
GSC 7.11
sh-ctrl
sh-ctrl + GSH
sh-ACADM1
sh-ACADM1 + GSH
Number of cells
Time (Days)
f
GSC 8.11 – 7 days
sh-ctrl
sh-ACADM1
Number of spheres/field
GSH
GSC 6.27 – 72h
sh-ctrl
sh-ACADM1
Number of spheres/field
GSH
GSC 6.27 – 7 days
sh-ctrl
sh-ACADM1
Number of spheres/field
GSH
g
****
MFI (CellRox)
(relative to sg-ctrl)
sg-ctrl
sg-ACADM1
sg-ctrl
sg-ACADM1
LPDS

Figure S8
